# Heterologous Surface Display Reveals Conserved Complement Inhibition and Functional Diversification of *Borrelia burgdorferi* Elp Proteins

**DOI:** 10.1101/2024.08.23.609448

**Authors:** Nathan Hill, Lara M. Matulina, Cameron MacIntyre, M. Amine Hassani, Sheila Thomas, Matteo Luban, Isabelle Ward, Amina Abdalla, John M. Leong, Brandon L. Garcia, Jacob E. Lemieux

## Abstract

Lyme disease is a tick-borne spirochetosis with diverse clinical manifestations. Phenotypic variation among *Borrelia burgdorferi* strains correlates with variable manifestations of Lyme disease in humans; this diversity is attributed in part to variation in surface-exposed lipoproteins, which are targets of the human antibody response and contribute to tissue adhesion, immune evasion, and other host interactions. Many *B. burgdorferi* lipoproteins are encoded as multi-copy gene families including the OspE/F-like leader peptide (Elp) protein family. To characterize Elp allelic variants, we adapted the *Pseudomonas syringae* ice nucleation protein (INP) system to present *B. burgdorferi* lipoproteins on the surface of *Escherichia coli*. We identified interactions with classical complement proteins and mapped binding regions in the *E. coli* system. We validated interactions using recombinant proteins and *B. burgdorferi* surface display. By assessing potential interactions with extracellular matrix components, we identified a novel interaction between Elp proteins and perlecan, a component of mammalian basement membranes, and revealed the bifunctional nature of Elps. Our findings reveal that Elps have undergone functional diversification while maintaining classical complement inhibition mediated by potent and conserved C1s binding and demonstrate that *E. coli* surface display offers an efficient, cost-effective, and relatively high throughput approach to characterize *B. burgdorferi* lipoproteins.

## Introduction

Lyme disease is the most common vector-borne disease in North America, with an estimated 476,000 cases in the United States per year (Kugeler *et al*., 2021). Case numbers are expected to increase across an expanding geographic range (Mead, 2015; Bouchard *et al*., 2015; Lantos *et al*., 2015). Lyme disease is clinically variable, with diverse manifestations that differ from person-to-person. Disease usually begins with a rash at the site of bacterial inoculation by tick bite; if untreated, the pathogen may disseminate to secondary sites including the heart, joints, and nervous system (Tan *et al*., 2021).

Lyme disease is caused by tick-borne spirochetes from *Borrelia burgdorferi* sensu lato, a phylogenetic subgroup which contains over twenty genospecies, including *B. burgdorferi* sensu stricto, *B. afzelii*, *B. garinii,* and *B. bavariensis,* the four genospecies that cause most human infections (Rudenko *et al*., 2011). Genetic variation in these species has been shown to influence the severity and clinical manifestations of Lyme disease in humans. For example, *B. garinii* and *B. bavariensis* are more likely to cause neuroborreliosis (van Dam *et al*., 1993; Balmelli and Piffaretti, 1995), whereas *B. afzelii* is associated with acrodermatitis (van Dam *et al*., 1993; Balmelli and Piffaretti, 1995), and *B. burgdorferi* sensu stricto is more likely than other genospecies to cause arthritis (Grillon *et al*., 2018). Among genotypes of *B. burgdorferi* sensu stricto, RST1 and OspC Type A are associated with higher rates of dissemination (Wormser *et al*., 1999; Dykhuizen *et al*., 2008; Wormser *et al*., 2008; Lemieux *et al*., 2023), increased rates of persistent arthritis (Jones *et al*., 2009), and greater severity (Strle *et al*., 2011). Thus, microbial variation strongly influences the clinical phenotype of Lyme disease.

Pathogenic spirochetes, including *B. burgdorferi,* encode an unusually large number of surface lipoproteins in their genomes, relative to other bacteria (Haake, 2000). Lipoproteins are major targets of human antibody response, but also play a critical role in virulence by preventing immune-mediated killing and promoting bacterial attachment to host tissues (reviewed in (Antonara *et al*., 2011; Steere *et al*., 2016; Lin *et al*., 2020a)). *B. burgdorferi* harbors ∼85 surface lipoproteins that are thought to influence strain-specific properties. Lipoproteins are encoded primarily by the plasmids that make up the *B*. *burgdorferi* genome; these plasmids are strain-variable, resulting in a variable lipoproteome with extensive variation among clinical isolates (Lemieux *et al*., 2023; Lemieux, 2024). *Borrelia* spp. encode over 1500 lipoprotein ortholog groups, most of which have not been experimentally characterized (Radolf *et al*., 2021).

A recent analysis of *B. burgdorferi* pangenomes revealed that linked clusters of accessory genome elements, predominantly encoded on plasmids and enriched in surface lipoproteins, correlated with rates of dissemination (Lemieux *et al*., 2023). The size of the lipoproteome correlated with the probability of dissemination, and strains that disseminate were found to have higher numbers of lipoproteins, an effect that was driven by multi-copy Mlp and Erp gene families, suggesting that members of these gene families may contribute to dissemination. Erps, which are OspEF-related proteins are categorized into three subfamilies: OspE-related Erps, OspF-related Erps, and OspE/F-like leader peptide (Elp) proteins (Akins Darrin R. *et al*., 1999). These proteins cluster within subfamilies and possess functional similarities. Multiple members of the OspE-related Erp family bind to Factor H, while several members of the OspF-related Erp family bind to glycosaminoglycans like heparan sulfate (Lin *et al*., 2015). The Elp family had lacked characterization for many of its members, until two members, ElpB and ElpQ, were shown to possess affinity for multiple complement components (Pereira *et al*., 2022). These proteins were shown to inhibit complement activation and subsequent killing, improving serum survival and bacterial fitness. *Borrelia* with increased ability to evade complement will likely cause more severe disease (reviewed in (Skare and Garcia, 2020)). Another Elp protein, ElpX, was shown to bind to laminin, an extracellular matrix component (Brissette *et al*., 2009); this binding function could improve bacterial adhesion and impact tissue tropism. Disparate binding functions within the same protein family pose interesting questions of conservation of binding affinity across the Elp protein family, as well as of potential bifunctionality for Elps. In addition to genetic and functional diversity within a gene family, which has been studied in only one or a few reference strains, the allelic diversity of these gene families has not been characterized. Thus, further understanding of the Elp family which has proven to have functional significance will improve understanding of Lyme disease manifestations, but the multiple copies of these gene families, presumed functional redundancy, and extensive allelic diversity have made multi-copy gene families difficult to study in *B. burgdorferi* and many other pathogens.

Previously, a *B. burgdorferi* overexpression library representing the B31 lipoproteome (Dowdell *et al*., 2017) was used to identify the ElpB/Q interactions; however extensive screening using *B. burgdorferi* surface display presents challenges. There is extensive microbial genetic diversity, including within the lipoproteome (Casjens *et al*., 2017; Lemieux *et al*., 2023; Akther *et al*., 2024). The Elp family alone possesses dozens of homologs (Akins Darrin R. *et al*., 1999; Pereira *et al*., 2022; Lemieux *et al*., 2023). It is challenging to generate *Borrelia* libraries of large size, as *B. burgdorferi* molecular genetics remains difficult. This spirochete grows slowly in *in vitro* culture, with a doubling time of 12-16 hours (Barbour, 1984; Norris *et al*., 1995).

Transformation efficiency is low, particularly among clinical strains, which possess restriction endonucleases (Samuels, 1995; Seshu *et al*., 2021). *B. burgdorferi* surface display also has the drawback of confounding, natively-expressed lipoproteins. Despite repetitive passaging, which leads to plasmid loss and reduction of native lipoproteins (Grimm *et al*., 2003), *B. burgdorferi* requires some plasmids, e.g. lp54 (Casjens, 1999) to survive under *in vitro* growth conditions. Lipoproteins encoded on essential plasmids can make interpretation of differential binding results difficult. Overall, *Borrelia*-based applications are limited by high cost, low throughput, and extended time required to generate libraries of transformants in *B. burgdorferi*. Efficient methods that overcome these experimental challenges are needed to characterize the biological roles within the extensive and diverse *B. burgdorferi* lipoproteome.

Heterologous display systems provide an alternative to display *Borrelia* lipoproteins and have been explored to bypass *Borrelia*-specific experimental challenges. Phage display libraries expanded the scope of *Borrelia* lipoprotein binding screens, leading to the identification of *B. burgdorferi* P66 (Coburn et al., 1999), an integrin ligand, as well as several other adhesins (Antonara et al., 2007). Initial attempts to use *Escherichia coli* to surface display *B. burgdorferi* lipoproteins were unsuccessful (Dunn et al., 1990), likely because of differences in the lipoprotein trafficking machinery. Robertson et al. modified *E. coli* to accomplish display of *B. burgdorferi* lipoproteins (Robertson et al., 2019). Their approach shows a degree of conservation of secretion mechanisms between *Borrelia* and *E. coli*; BBA57, with the native signal sequence intact, was translocated to the *E. coli* outer membrane and properly folded. Proteins involved in export with homologs in *Borrelia* include LolA (a periplasmic shuttle protein) (Sutcliffe et al., 2012), as well as BamA and BamB (which help form the β-barrel assembly machine) (Stubenrauch et al., 2016). However, not all lipoproteins were surface expressed when using the native signal sequences, indicating interspecies commonality in secretion mechanisms has limits. These challenges showed the necessity of developing a new system to screen *B. burgdorferi* lipoproteins.

We sought to define a new approach that retains features of *B. burgdorferi* surface display (Dowdell *et al*., 2017; Pereira *et al*., 2022) while reducing time, cost, and complexity. Multiple methods allow *E. coli* to express heterologous proteins on the bacterial surface, usually involving fusion to a surface localization sequence and a membrane tethering motif. One of these methods, developed by Jung et al., fuses a heterologous protein domain to the C-terminus of the N-terminus of the *Pseudomonas syringae* Ice Nucleation Protein (INPN) (Jung *et al*., 1998), which is sufficient to localize the fusion protein to the *E. coli* outer membrane. In this study, we adapted the INPN- based heterologous surface display system to present *B. burgdorferi* lipoproteins, including those from the Elp protein family, on the surface of *E. coli*. We demonstrated that this system serves as an efficient tool to characterize novel *B. burgdorferi* lipoprotein interactions and identified conserved complement inhibitory properties and functional diversification of the Elp family.

## Results

### Establishing an E. coli INPN-based surface display system for B. burgdorferi lipoproteins

In an effort to develop tools to enable the efficient study of lipoproteome variation, such as lipoprotein allelic variants from recent pangenome analyses (Lemieux *et al*., 2023), we pursued surface display in *E. coli*, a host with high transformation efficiency, short growth time, and minimal biosafety requirements. To adapt the INPN- based surface display for *B. burgdorferi*, we engineered *E. coli* to express mature *B. burgdorferi* lipoprotein, i.e, lacking the native lipobox sequences that are cleaved upon translocation. Because the machinery for trafficking *B. burgdorferi* lipoproteins to the cell surface is absent in *E. coli,* we included the non-native elements of *pelB* signal sequence (derived from *Erwinia carotovora*) (Lei *et al*., 1987) and INPN linker at the N-terminus of the fusion protein. We also included a C-terminal His_6_-tag for antibody-mediated detection. For initial testing, we generated an *E. coli* strain that produces INPN alone (INPN) and INPN fused to the mature portion of *B. burgdorferi* lipoproteins CspA, which binds the complement regulator factor H (Kraiczy *et al*., 2001). Flow cytometric analysis of the *E. coli* transformant producing a CspA-INPN fusion (CspA), after staining with anti-His_6_ antibody revealed that surface expression, which was assessed in comparison to the same strain without antibody treatment, was higher after overnight growth than earlier time points (Figs. S1A, S1B), an effect we attribute to the de-repression of the T7 promoter during extended growth at 37°C (Grossman *et al*., 1998).

We extended our analysis to include an ElpQ-INPN fusion (ElpQ); ElpQ binds complement C1, C1r, and C1s (Pereira *et al*., 2022). Flow cytometric analysis of ElpQ and CspA strains revealed that 96% of cells produce surface-localized ElpQ above the background level, compared to 50% of CspA (Fig. 1A). Fluorescence microscopy performed after staining the two strains with anti-His_6_ antibody confirmed the presence of CspA and ElpQ on the surface of *E. coli*, again with an apparent higher degree of surface expression for ElpQ compared to CspA (Fig. 1B). We generated strains for multiple *B. burgdorferi* lipoproteins, including several B31 Elps (e.g. ElpQ, ElpB, and ElpX), and found high levels of surface expression (Fig. 1C). These experiments confirmed that structurally diverse *B. burgdorferi* lipoproteins, and Elps specifically, were surface expressed through the INPN system.

**Figure 1:**
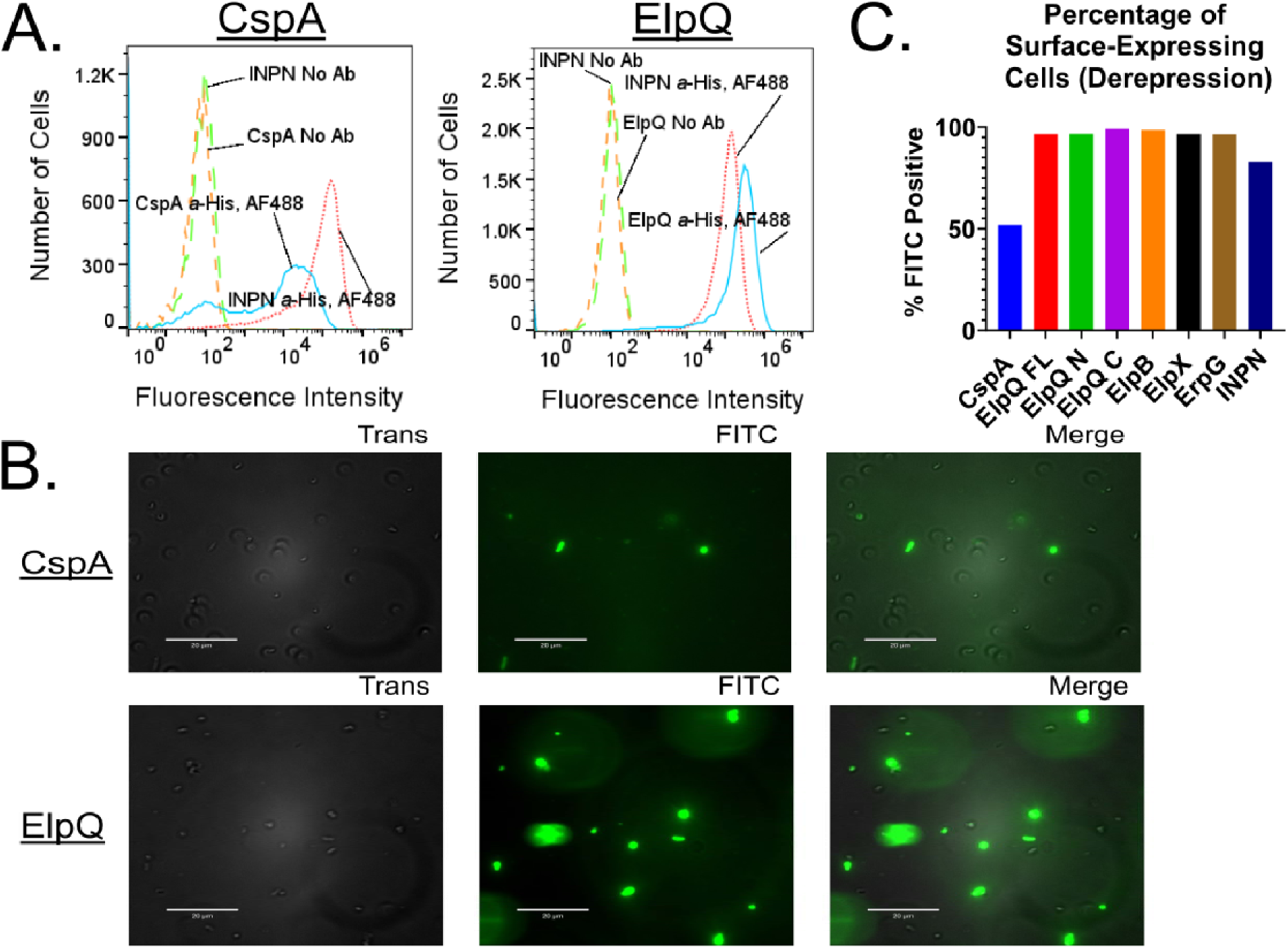
*Borrelia* ElpQ and CspA can be expressed on the surface of *E. coli* when fused to INPN: (A). 1×10^8^ cells per condition were collected from uninduced cultures of E. coli expressing INPN, CspA-INPN, or ElpQ-INPN grown to OD_600_≈1.5 and washed. Cells were blocked for 2 hr with 3% BSA and incubated overnight with a rabbit anti-His antibody. Cells were washed again and incubated for 1 hr with anti-rabbit AF488 antibody. Cells were fixed with 4% formaldehyde and 30000 events were analyzed on a BD Accuri™ C6 Flow Cytometer. (B). 1×10^8^ cells per condition were collected and treated as described above. Cells were observed at 100x magnification using an Echo Revolution through the FITC channel. (C). Percentage of FITC positivity for various strains grown overnight and detected through flow cytometry using anti-His and anti- rabbit AF488 antibodies.

### Heterologously displayed B. burgdorferi lipoproteins are functional

We used far western blotting to validate the interaction of *B. burgdorferi* proteins produced on the surface of *E. coli* with their known binding partners. Previously, when *B. burgdorferi* cultures overexpressing lipoproteins of interest were lysed and separated by SDS-PAGE, they were able to recognize their known binding targets through far western blotting (Pereira et al., 2022; McDowell et al., 2006). Incubation of the SDS-PAGE-separated proteins from *E. coli* producing surface-localized ElpQ with human complement C1s and an anti-C1s antibody resulted in a specific band roughly corresponding to the size of the ElpQ-INPN fusion (62.6 kDa), whereas no corresponding band was observed for the CspA or INPN negative control strains (Fig. 2A). Similarly, assessment of *E. coli* expressing CspA, probed with FH and an anti-FH antibody resulted in a band at roughly the expected size (51.6 kDa), while the other strains, ElpQ and INPN, did not produce bands (Fig. S2A). These western blotting results confirm that multiple *Borrelia* lipoproteins interact in the same way as natively-expressed *B. burgdorferi* lipoproteins when surfaced-expressed on *E. coli*.

**Figure 2:**
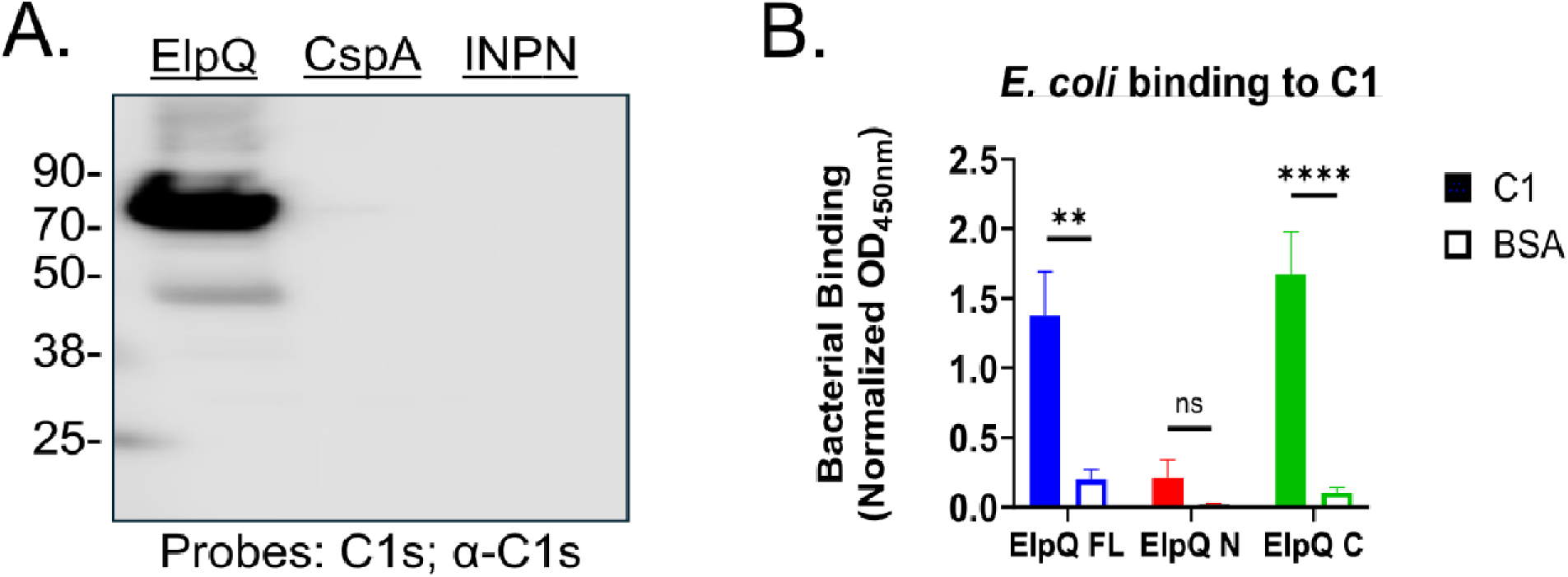
*Borrelia* ElpQ retains ligand-specific binding activity upon ectopic expression on the surface of *E. coli*: (A). Far western blotting was performed as described (Methods), using 2 µg/mL C1s. Binding was detected using mouse anti-C1s antibody and anti-mouse HRP-conjugated antibody. (B.) Strains expressing ElpQ-INPN, ElpQ N-INPN, and ElpQ C-INPN were grown overnight with no induction as previously described. 8×10^6^ cells were collected from cultures to add in quadruplicate to C1 and BSA-coated wells (along with PBS). Cells were centrifuged to encourage cell-well contact, and allowed to incubate for 1 hr. Afterwards non-adherent cells were washed away as before, and adherent cells quantified as before. ElpQ and ElpQ mutant binding was assessed in triplicate experiments (n=3). Error bars indicate SEM. ****P < 0.0001; *P < 0.05; ns, not significant using Student’s t test to compare mean values.

We performed whole cell ELISA, a high throughput screening method, to evaluate the binding of *E. coli*-tethered *B. burgdorferi* lipoproteins in their natively folded state without lysis. We compared the specific binding for surface-expressed proteins to the non-specific binding to BSA, the negative control. A recent study (Garrigues *et al*., 2022) determined an N-terminal portion of ElpQ (aa 19-216) lacked complement inhibitory activity, whereas a C-terminal portion of ElpQ (aa 168-343), inhibited complement to a similar level as full-length ElpQ. We assessed whether similar truncation mutants show similar results in INPN-based *E. coli* surface display. We generated three constructs, the full-length ElpQ lacking the native signal sequence (aa 19-343), along with N-terminal (aa 19-206) and C-terminal (aa 168-343) truncation constructs that were termed ElpQ FL, ElpQ N, and ElpQ C, respectively. These constructs were assessed for C1 binding using *E. coli* whole cell ELISA. In agreement with previous findings, the ElpQ FL and ElpQ C showed a significant binding to C1 (Fig. 2B). No significant binding to C1 was observed for ElpQ N, which served as the negative control. We also performed whole cell ELISA to evaluate binding of CspA to FH. CspA bound to FH significantly, but with low overall OD_450_ values (Fig. S2B). We then grew CspA under induction conditions and, despite lowered surface expression (Fig. S2C), noted significant binding with higher OD_450_ values in a slightly modified whole cell ELISA (Fig. S2D). Our results provide evidence that *B. burgdorferi* lipoproteins heterologously expressed on the surface of *E. coli* using INPN-based surface display maintain their binding behavior similarly to those natively expressed on *B. burgdorferi*.

### E. coli Strains Expressing non-B31 Elp Homologs Reveal Novel Binding to Complement Components

We assessed four proteins which are present in the *B. burgdorferi* pangenome but are absent in *B. burgdorferi* B31. These proteins share common residues with each other and other B31 Elps to differing degrees (Fig. S3). We labeled these genes in the order in which they were generated for study, BG036, BG044, BG032, and BG038. The closest matches to any named *Borrelia* and named B31 protein can be found in Supplemental Table 1 (Table S1). Phylogenetic analysis across strains (Fig. S4) indicated that BG032 is found in RST3/OspC type D, E, G and T strains, whereas BG044 is found predominantly in RST2/OspC type K and H strains. In contrast, BG036 is more widespread, found in RST1/OspC type B, RST2/OspC types I and K, and RST3 /OspC type I strains. BG038 was present in only a few clinical isolates.

These candidate genes showed homology to B31 ElpB/Q, which previous studies found bind to C1s, blocking the cleavage of C2 and C4 and inhibiting the initiation of the classical pathway. (Pereira *et al*., 2022; Garrigues *et al*., 2022) To test if this function was conserved in Elps beyond ElpB/Q, we generated strains expressing all four candidate proteins (i.e. BG036, BG044, BG032, and BG038). We expressed each non- B31 Elp homolog using this *E. coli* system under induction growth conditions based on the improved detection of CspA binding. As expected, this led to lower, and more variable amounts of surface expression (Fig. 3A). We screened the four candidates and the positive control B31 ElpQ using whole cell ELISA against human C1s. In all four candidate lipoproteins and the positive control, we detected significant binding to C1s in the *E. coli* screen (Fig. 3B). For BG032, we noted a high “background” of BSA binding, however, the higher overall binding signal for C1s suggested an interaction. Our results demonstrate that, like B31 ElpQ, all four lipoprotein candidates bind to human complement C1s protein, suggesting conserved functional properties of these lipoproteins in *B. burgdorferi*. Our results also validated the use of the induction growth conditions in detecting Elp interactions.

**Figure 3:**
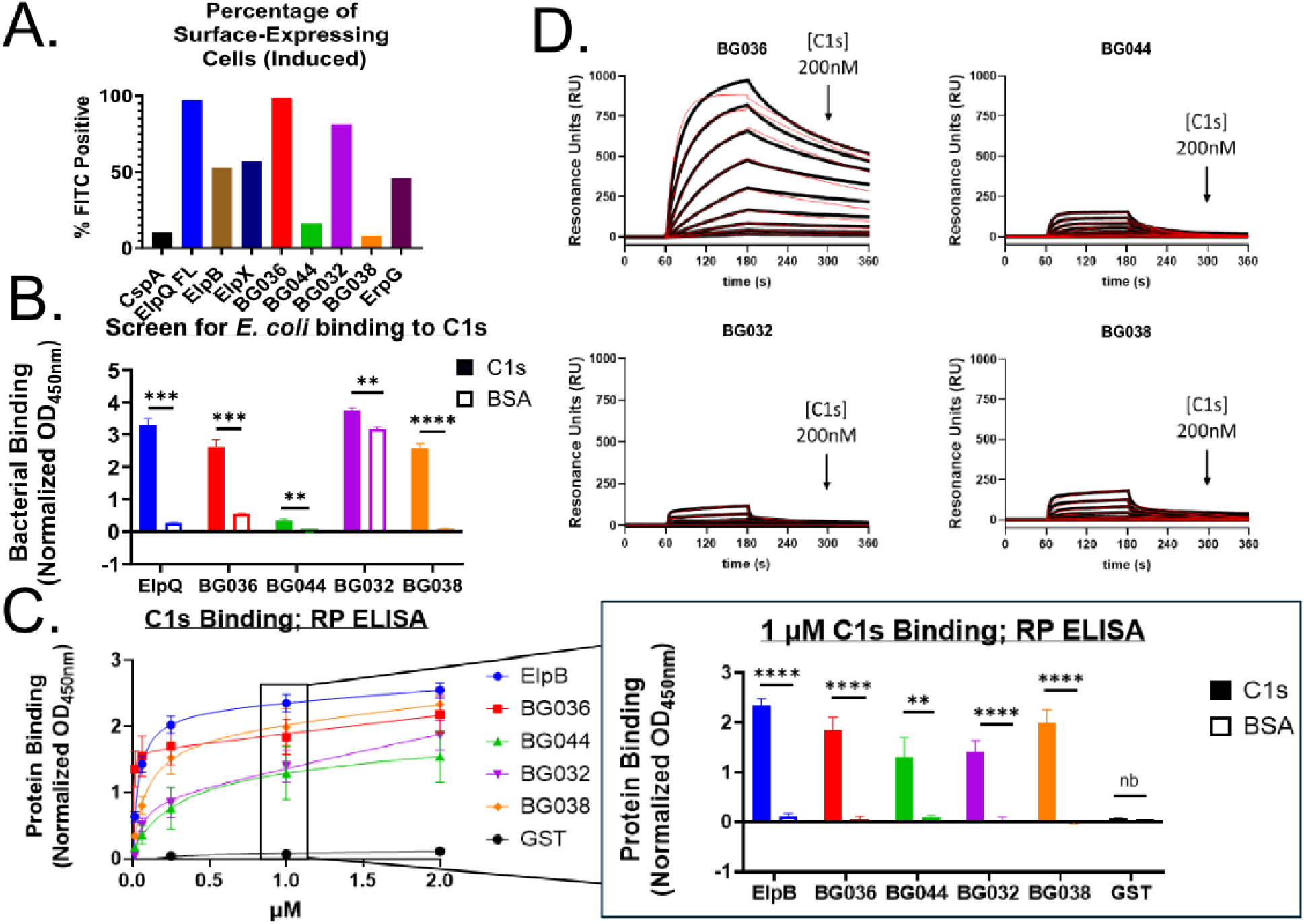
Conserved C1s Binding Function Among Elp Homologs: (A.) Percentage of FITC positivity for various strains grown under the induction condition and analyzed through flow cytometry using rabbit anti-His and anti-rabbit AF488 antibodies. Positivity determined as previously described. (B). Strains expressing ElpQ-INPN and the four selected Elp homologs linked to INP were grown to ∼0.6 OD_600_ and induced for 18 hr as previously described. 8×10^8^ cells were collected to add in quadruplicate to C1s and BSA-coated wells. Whole cell ELISA was performed as previously described in a single experiment (n=1). Error bars indicate SEM. ****P < 0.0001; *P < 0.05; ns, not significant using Student’s t test to compare mean values. (C). ElpB and Elp homolog recombinant proteins, along with GST, were added in a dilution series starting at a concentration of 2 μM to wells coated with C1s and BSA. ELISA was carried out as previously described with protein binding quantified through anti-GST and anti-GST HRP incubation. Each recombinant protein was assessed at least in triplicate (n=3). The highest BSA background mean value assessed was 0.256 OD_450_. Error bars indicate SEM. ****P < 0.0001; *P < 0.05; ns, not significant using Student’s t test to compare mean values. (D). Each Elp homolog was immobilized on an SPR sensor chip. Purified activated human C1s was injected over each biosensor in a two-fold dilution series ranging from 0 to 200 nM. Sensorgrams from a representative injection series are shown. Each injection series was performed in triplicate (n=3). Sensorgrams were fit to a 1:1 kinetic model of binding (Langmuir). The raw sensorgrams are shown in black and the kinetic fits in red.

Based on these results, we generated recombinant proteins of these lipoprotein candidates and B31 ElpB, a positive control, and tested their binding to C1s in recombinant protein ELISA. We used purified GST tag as a negative control and compared all specific signals to non-specific binding to BSA. ElpB showed the strongest binding to C1s, followed by BG036, BG038, BG032, and BG044 (Fig. 3C). The recombinant BG032 did not show strong binding affinity to BSA, indicating the specificity of the interaction between BG032 and C1s. We next measured equilibrium dissociation constants (*K*_D_) of ‘tag-free’ proteins using surface plasmon resonance (SPR) (Fig. 3D). In the SPR experiment, binding was strongest for BG036, followed by BG044, BG038, and BG032, corroborating the whole cell ELISA results. Our findings using both methods confirm the interaction of these Elp lipoprotein candidates with human complement C1s, and further validate the approach of screening uncharacterized proteins with INPN-based surface display, although with some differences in relative binding affinities across methods.

It has been previously shown that ElpB/Q bind with high affinity to C1r (Pereira *et al*., 2022). However, neither lipoprotein was found to block C1r from cleaving its downstream complement cascade substrate, C1s proenzyme. We then investigated whether C1r binding affinity was conserved among these four lipoprotein candidates using *E. coli* whole cell ELISA. B31 ElpQ, BG032, and BG038 showed similar levels of specific and nonspecific binding compared to the C1s screen, whereas BG036 and BG044 showed a weaker interaction and no interaction, respectively (Fig. 4A). BG032 and BG038 were assessed through recombinant protein ELISA in triplicate, while BG036 and BG044 were assessed in a single recombinant protein ELISA. The protein candidate BG032 exhibited a high background signal. Recombinant protein ELISA using GST-tagged proteins showed low levels of binding to C1r for all four candidate lipoproteins (Fig. 4B) compared to their binding for C1s. All proteins tested showed noticeably lower binding affinity than BBK32, the positive control. Consistent with these results, we could not detect binding to C1r for any of the non-B31 homologs using SPR at concentrations up to 200 nM using ‘tag-free’ proteins (Fig. 4C). These results clearly indicate that interactions between the four Elp lipoprotein candidates and C1r, while still detectable, are of lower affinity compared to their interactions with C1s. In addition, this data showed that *E. coli* whole cell ELISA results cannot be used directly to show binding affinity. BG032 and BG038 showed similar binding signals for C1r and C1s, despite both possessing higher affinity for C1s. Multiplicity of proteins on the bacterial surface can produce a high signal in the absence of high affinity, showing the importance of orthogonal assays in assessing binding affinity. However, for identifying interactions, the *E. coli* whole cell ELISA proved its utility.

**Figure 4:**
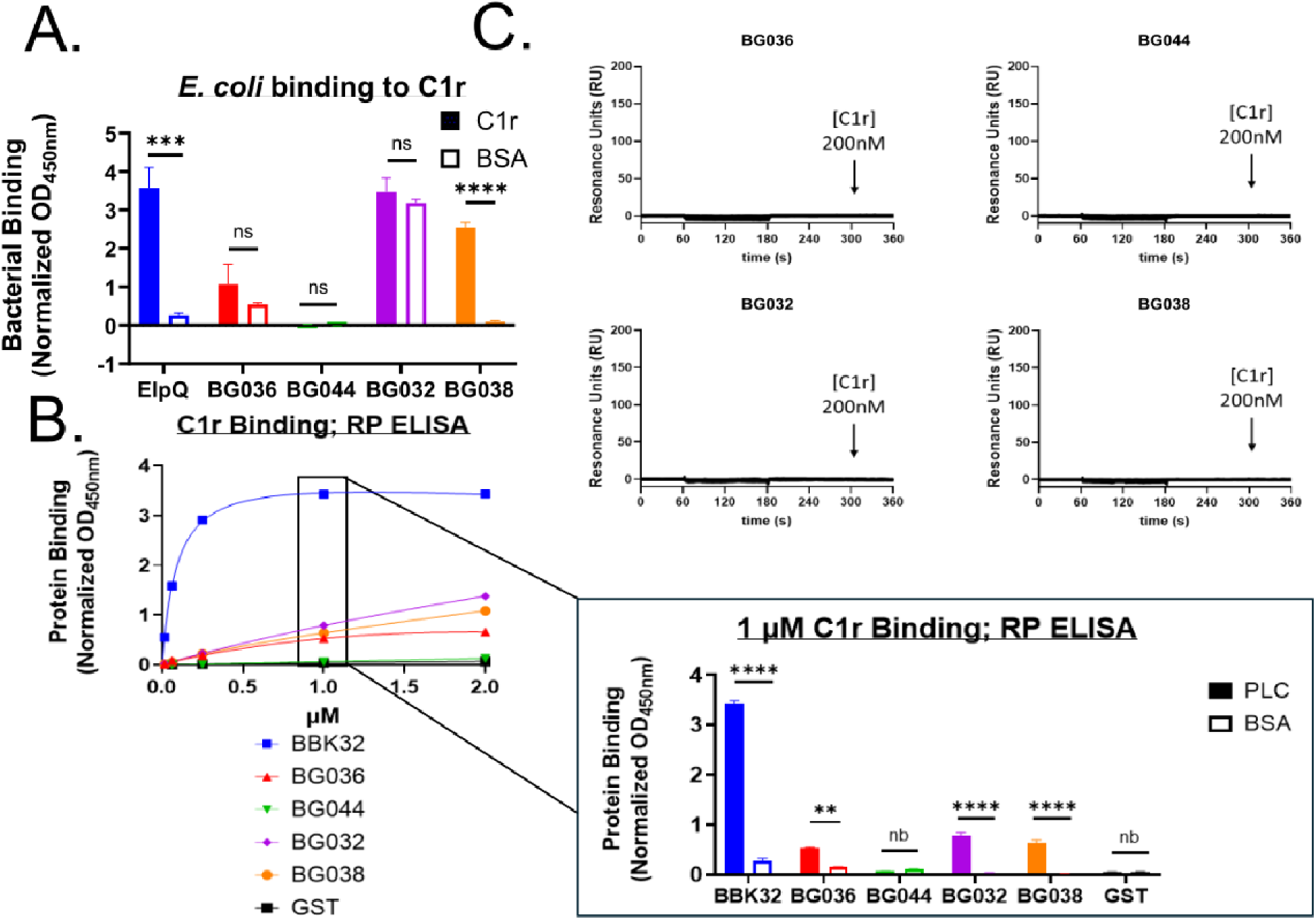
Variable C1r Binding Among Elp Homologs: (A). Strains expressing ElpQ- INPN and the four selected Elp homologs linked to INP were grown to ∼0.6 OD_600_ and induced for 18 hr as previously described. 8×10^8^ cells were collected to add in quadruplicate to C1r and BSA-coated wells. Whole cell ELISA was performed as previously described in a single experiment (n=1). Error bars indicate SEM. ****P < 0.0001; *P < 0.05; ns, not significant using Student’s t test to compare mean values. (B). BBK32 and Elp homolog recombinant proteins, along with GST, were added in a dilution series starting at a concentration of 2 μM to wells coated with C1r and BSA. ELISA was carried out as previously described with protein binding quantified through anti-GST and anti-GST HRP incubation. BG032 and BG038 recombinant proteins were assessed at least in triplicate (n=3). BG036 and BG044 recombinant proteins were assessed with one replicate (n=1). Error bars indicate SEM. ****P < 0.0001; *P < 0.05; ns, not significant using Student’s t test to compare mean values. The highest BSA background mean value assessed was 0.256 OD_450_. (C). Each Elp homolog was immobilized on an SPR sensor chip. Purified activated human C1s was injected over each biosensor in a two-fold dilution series ranging from 0 to 200 nM. Sensorgrams from a representative injection series are shown. Each injection series was performed in triplicate (n=3).

### Non-B31 Elps are Complement Inhibitors

To characterize the functional implications of this binding pattern, we used a serum-based classical pathway activation ELISA and measured C4b deposition, which is indicative of complement activation. All four candidate proteins displayed dose-dependent inhibition of complement activation; BG036 and BG044, which showed strong binding to C1s in multiple binding assays, exhibited potent complement inhibition with IC_50_ values of 89.8 nM and 139.7 nM, respectively (Fig. 5). In contrast, the other two candidates Elps, BG032 and BG038, inactivated complement to a lesser extent than BG036 and BG044, with an IC_50_ roughly one order of magnitude higher (Table S4). Our data demonstrate that these proteins show differential levels of complement inhibition and suggest a role for all in immune evasion.

**Figure 5:**
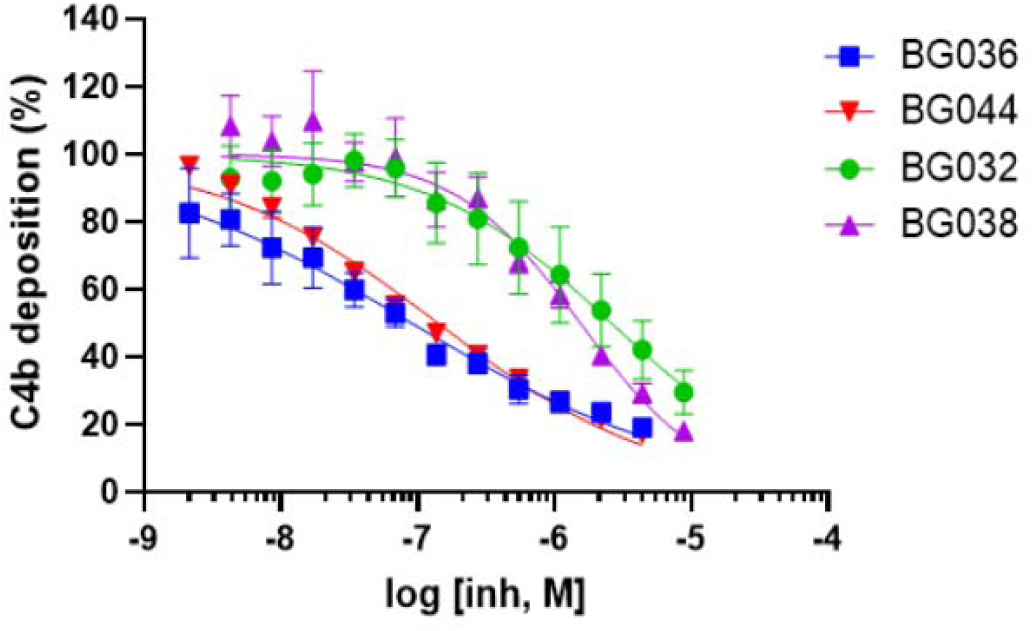
Elp homologs inhibit the classical pathway of complement: Classical-pathway specific complement activation was measured using an ELISA-based assay. C4b deposition was used as a marker for activation and followed in the presence of a concentration series of each Elp homolog. BG036 is depicted in blue, BG044, red, BG032, green and BG038, purple (n=4). Error bars depict SD. IC_50_ values were obtained by nonlinear regression analysis using a four-parameter variable slope fit, constraining the bottom and top values to 0 and 100, respectively.

### E. coli Strains Expressing non-B31 Elp Homologs Suggest Potential Binding to Perlecan

We next used *E. coli* INPN-based surface display to identify potential novel ligands for Elp proteins. We broadened our screens to include components of the extracellular matrix (ECM), which are known to interact with *B. burgdorferi* lipoproteins. B31 ElpX, a member of the Elp family, binds to laminin, an ECM component (Brissette *et al*., 2009). We generated an ElpX strain and measured levels of surface expression under the derepression and induction growth conditions (Figs. S6A, S6C). We performed whole cell ELISA using ElpX grown under both conditions; the expected binding for ElpX was not observed for the derepression growth conditions (Fig. S6B) but was observed for the induction growth condition (Fig. S6D), further validating the use of that growth condition for diverse proteins and ligands. To assess whether laminin binding affinity was conserved among the candidate proteins, we performed *E. coli* whole cell ELISA, however the candidate proteins did not bind to laminin (data not shown).

We then screened perlecan, a proteoglycan possessing laminin-like regions. Perlecan harbors a core protein size of 435 kDa, spread across multiple domains, as well as heparan sulfate and chondroitin sulfate glycosaminoglycan side chains. Elps have not previously been shown to bind glycosaminoglycans; however, to control for heparin-mediated perlecan interactions, we generated a strain expressing an ErpG- INPN fusion (ErpG), as ErpG is a known heparin binder (Lin *et al*., 2015). Far western blots revealed that heparin binding was not sufficient to produce detectable perlecan binding, evidenced by the lack of a band for ErpG (Fig 6A). However, it was notable that two Elps, ElpQ and ElpB, that were included in the screen produced bands at their expected and experimentally-determined sizes (Fig. S7).

**Figure 6:**
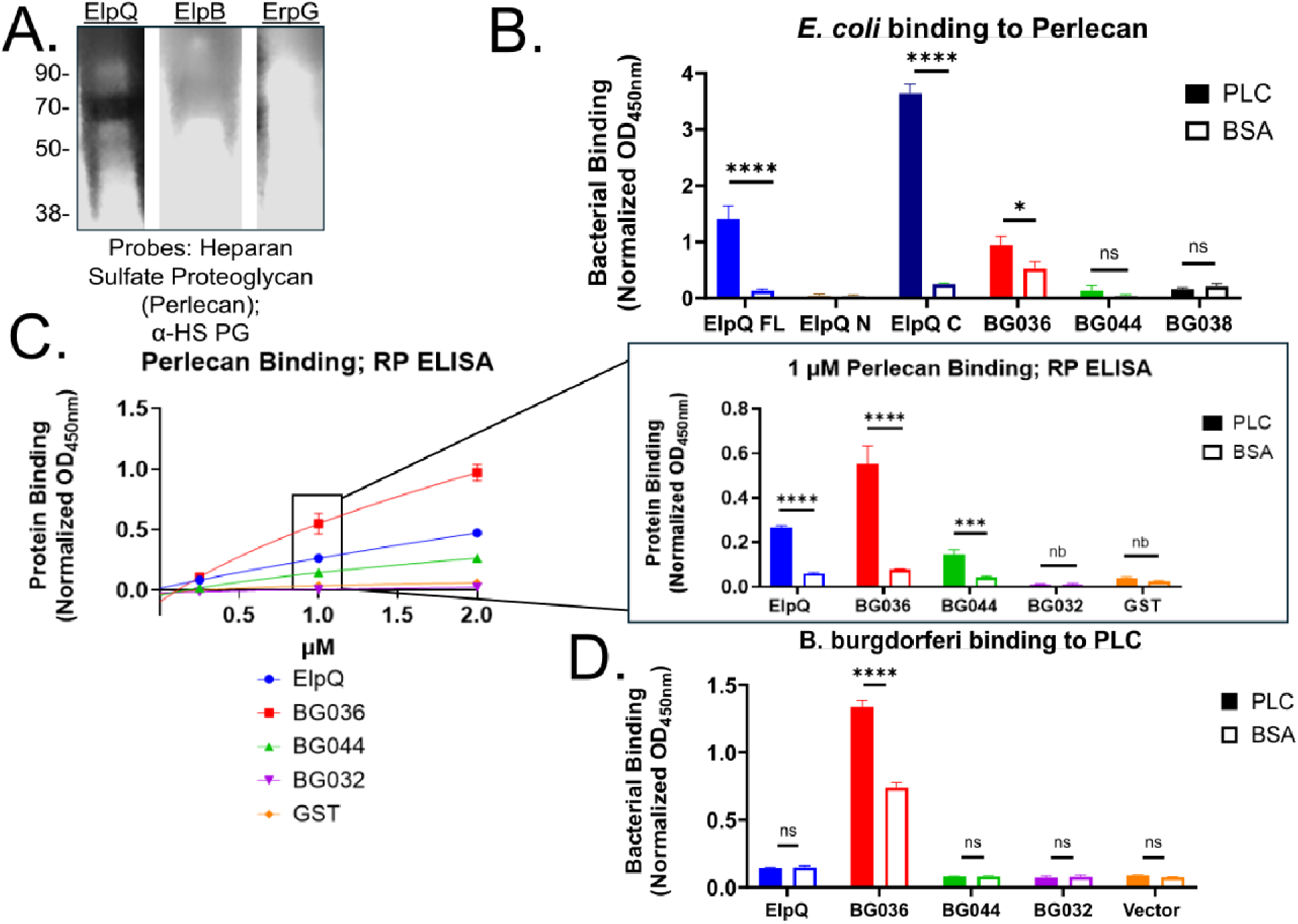
Variable Perlecan Binding Function Among Elp Homologs: (A). Far Western was performed as previously described. A membrane was incubated overnight with 2 µg/mL perlecan and detected with anti-HSPG (Sigma Aldrich MABT12) and anti-mouse HRP antibodies. (B). Strains expressing ElpQ-INPN, ElpQ N-INP, ElpQ C-INPN, and three Elp homologs linked to INP were grown under induction conditions. 8×10^8^ cells were collected to add in quadruplicate to perlecan and BSA-coated wells. Whole cell ELISA was performed as previously described in a single experiment (n=1). Follow-up whole cell ELISA was performed using strains that showed binding above background after the initial screen. Followed-up strains (ElpQ FL, ElpQ N, BG036, BG044) were assessed through quadruplicate experiments (n=4). Error bars indicate SEM. ****P < 0.0001; *P < 0.05; ns, not significant using Student’s t test to compare mean values. (C). Elp homolog recombinant proteins, along with GST, were added in a dilution series starting at a concentration of 2 μM to wells coated with perlecan and BSA. ELISA was carried out as previously described with protein binding quantified through anti-GST and anti-GST HRP incubation. Binding was assessed through triplicate experiments (n=3). The highest BSA background mean value assessed was 0.119 OD_450_. Error bars indicate SEM. ****P < 0.0001; *P < 0.05; ns, not significant using Student’s t test to compare mean values. (D). *Borrelia burgdorferi* B31 strains expressing BG036, BG044, BG032, and an empty pBSV2 vector were assessed through whole cell ELISA. 1.6×10^7^ cells were collected to add in quadruplicate to perlecan and BSA-coated wells. PBS was added to coated wells as previously described. Cells were centrifuged to encourage contact with ligand-coated wells and incubated for 2 hr. ELISA was carried out as described (Methods). Binding was assessed through duplicate experiments (n=2). Error bars indicate SEM. ****P < 0.0001; *P < 0.05; ns, not significant using Student’s t test to compare mean values.

We performed *E. coli* whole cell ELISA on B31 ElpQ, as well as the four Elp candidate proteins. There was a level of binding above the background for ElpQ FL, as well as BG036 and BG044 (Fig. 6B). Using the previously generated ElpQ deletion strains, we observed similar binding for ElpQ FL and ElpQ C. This, combined with the result for ElpQ N which showed no binding, provided an indication that the binding site for ElpQ to perlecan is likely at the C-terminal half of the protein. BG032 was not screened using the *E. coli* system due to the previous high level of binding to BSA in the INPN fusion form. BG038 did not show binding to perlecan and was not characterized further. We repeated the whole cell ELISA for the identified targets ElpQ, BG036, and BG044, as well as ElpQ N, which served as a negative control. After further confirmatory screening, we found that the ElpQ and BG036 bound significantly, while ElpQ N did not bind as expected. BG044 binding, while not significant, was high enough to merit further characterization.

We assessed the ElpQ FL, BG036, BG044, and the uncharacterized BG032 through recombinant protein ELISA. ElpQ, BG036, and BG044 bound at low levels (Fig. 6C) and these interactions appeared weak (Table S7). However, there is a clear increase in binding for these three proteins that is especially visible at the 1 μM concentration. There was little to no binding for BG032. The GST tag alone produced no binding as expected. We attempted to characterize these interactions by SPR but detected very low binding responses to perlecan with concentrations up to 200 nM (data not shown), an observation which may be related to the low overall perlecan affinity.

We then tested ElpQ, BG036, and BG044’s affinity in the native *Borrelia* host through *B*. *burgdorferi* whole cell ELISA. *B. burgdorferi* expressing BG036 had significantly higher binding to perlecan-coated wells compared to the BSA control (Fig. 6D). BG044 did not bind perlecan in this assay. ElpQ also did not bind to perlecan, though difficulties in consistently expressing the protein to high levels in recombinant *Borrelia* complicate this finding (Fig. S8). A vector control carrying only the pBSV2 vector in the B314 background was used as a negative control. BG032, which was not expected to bind to perlecan, was used as an additional negative control and did not bind.

There was heterogeneity in perlecan binding across assay methods and protein candidates. For BG036, an interaction was observed through *E. coli* whole cell ELISA, recombinant protein ELISA, and *B. burgdorferi* whole cell ELISA, but not SPR. For ElpQ and BG044, an interaction with perlecan was observed by *E. coli* whole cell ELISA and recombinant protein ELISA. An interaction was not observed by SPR or *B. burgdorferi* whole cell ELISA for either BG044 or ElpQ, although for the ElpQ strain, the lipoprotein displayed poorly and was observed to possess very low fluorescence values on *B. burgdorferi* (Fig. S8). BG032 was not assessed through *E. coli* whole cell ELISA, but did not show binding in recombinant protein ELISA, SPR, or *B. burgdorferi* whole cell ELISA. In contrast to other homologs, BG038 did not bind to perlecan through *E. coli* whole cell ELISA and as result was not assessed by recombinant protein, SPR, or *B. burgdorferi* whole cell ELISA.

## Discussion

We have shown here that diverse *B. burgdorferi* Elp proteins inhibit complement through conserved C1 binding. Potent inhibition of complement was observed even for allelic variants that without detectable C1r binding, indicating that C1s binding is sufficient for C1-mediated inhibition. This extends a previous observation that C1s affinity was greater than C1r for ElpB/Q (Pereira *et al*., 2022). In addition, the two Elp homologs with the most potent inhibition of C4b deposition were those with the strongest C1s binding according to SPR (Fig 3D). The sufficiency of C1s binding for complement inhibition and the correlation of C1s affinity with complement inhibition indicate that C1s binding is the dominant mechanism by which Elp-mediated inhibition of classical complement is achieved.

Characterizing allelic variants of Elp proteins also revealed functional diversification. Complement inhibition was a fixture across all Elp allelic variants, but the potency of inhibition varied, with the the most potent inhibition observed for BG036 and BG044. More notably, we observed a capacity for additional binding functions among family members: BG036 binds perlecan, while ElpQ is a potential interactor (Fig. 6); BG036 and ElpQ bind tightly to C1s and potently inhibit complement. Perlecan is a proteoglycan that is widely expressed in basement membranes (Hayes *et al*., 2022) and has been identified as a binding target promoting adhesion in other pathogens (Chen and Hazlett, 2000). The full functional significance of perlecan binding is unknown and will be important to assess in future experiments. Bifunctionality is common among *B. burgdorferi* lipoproteins, including those with complement inhibitory properties. For example, BBK32 binds C1r and inhibits complement (Garcia *et al*., 2016), while also binding fibronectin (Probert and Johnson, 1998) and dermatan sulfate (Fischer *et al*., 2006). Similarly, OspC binds to C4b (Caine *et al*., 2017) and fibronectin (Lin *et al*., 2020b). Recombinant protein assays suggest that the binding affinity is weak, but strengthening of weak interactions when proteins were displayed on a bacterial surface was also observed to a lesser extent for BSA, a highly negatively charged ligand (Figs. 6B, 6D). Promiscuous, weak interactions to host proteins based on charge, hydrophobicity, or other biochemical properties, may produce avidity effects that allow for specific interaction with perlecan in the absence of high affinity, providing a mechanism for binding to diverse host tissues. This is a mechanism that is used by other bacteria (Alvarez-Dominguez and Vazquez-Boland, n.d.), as well as other microbes such as viruses (Spillmann, 2001) to promote invasion. These low affinity interactions involving Elps could play a similar role in *Borrelia* infection, while maintaining critical complement inhibitory functions.

Our use of *E. coli* for this purpose extends prior work on *E. coli* heterologous surface expression (Robertson *et al*., 2019). A powerful advantage of using *E. coli* to display lipoproteins is the relative ease, low cost, and short timeframe for generating and characterizing mutants. All strains grow in Luria Broth (LB), which is accessible, inexpensive, and consistent across laboratories. Strains, once expressed through cloning, can be transformed, stocked, and screened for surface expression within 4 days. Functional screening can begin immediately after, with *E. coli* whole cell ELISA taking only 3 days to perform, starting from a frozen stock. Addition to and regeneration of libraries is relatively trivial. The entire process also takes place at BSL1, eliminating biosafety concerns. When working with *Borrelia*, the clone production and validation process takes weeks, requires more hands-on time, and clone isolation is more complex due to the need for limiting dilution or plating on semi-solid agar. Like *B. burgdorferi*-based surface display, our system also maintains the biological context of surface-exposed lipoproteins, by expressing them on a cell surface. A limitation of the system is the variable levels of expression for certain proteins, a potential confounding factor in assessing strengths of interactions based solely on *E. coli* whole cell ELISA results. Nevertheless, variable surface abundance was also an issue for *B. burgdorferi* surface display and is, therefore, not unique to *E. coli*. Recombinant protein assays tended to have lower background and greater flexibility, but recombinant protein is more expensive and time-consuming to produce, and binding occurs in a more artificial context that may miss important effects related to a cell surface. Thus, no single approach is optimal; rather, a toolkit of approaches affords different advantages and drawbacks that make them suited for various purposes. *E. coli* surface display seems particularly well suited for situations in which lipoproteins express well on the surface of *E. coli* and it is of interest to investigate the binding properties of large number of variants, for example allelic variants or mutational scanning experiments.Overall, our data show that Functional diversification of immune evasion and host adhesion could contribute to strain-mediated variability of human clinical disease and includes strain-mediated differences in dissemination (Wormser *et al*., 1999; Jones *et al*., 2006; Dykhuizen *et al*., 2008; Wormser *et al*., 2008; Lemieux *et al*., 2023) and tissue tropism (Jones *et al*., 2009). BG036 and BG044 are present in OspC types H and K, strains with a high probability of invasion (Lemieux *et al*., 2023). Perlecan binding affinity, which seems to be strongest in BG036, potentially gives strains an advantage in adhesion, improving colonization and fitness. These observations suggest the importance of correlating various functional properties of infecting spirochetal strains with clinical manifestations in larger-scale translational studies, likely at the site of disease (e.g. within an erythema migrans lesion, a joint, or in the nervous system) as a future avenue of investigation to assess the relative importance of these bacterial properties in the context of human infection.

In summary, our analysis of variation among Elp homologs reveals the conserved nature of classical pathway complement inhibition across naturally-occuring *B. burgdorferi* strains. Genetic and functional data indicate that this effect is largely mediated by C1s binding. In characterizing Elp homologs, we further developed *E. coli* surface display as a useful tool to characterize the extensive strain- and allelic variability among surface lipoproteins of *B. burgdorferi*. The lipoproteome diversity of *Borrelia* lipoproteins, which number approximately ∼85 in a single Lyme disease strain (Fraser *et al*., 1997; Casjens *et al*., 2000; Dowdell *et al*., 2017) and exceed 1500 across the *Borreliaceae (Lemieux, 2024)*. Given the importance of surface lipoproteins across pathogenic spirochetes, understanding the functional consequences of surface lipoprotein variation is critical for reducing the growing burden of disease attributable to this class of organisms.

## Experimental Procedures

### E. coli plasmid generation

Plasmids are generated as previously described (Fan *et al*., 2011). pETN is the most basic strain experimentally examined, expressing only the His-tagged INPN linker. pETN was generated by restriction cloning using EcoRI and HindIII to clone a codon-optimized version of the N-terminal portion of the INP gene from *Pseudomonas syringae* into pET22b (Novagen). This INPN gene fragment was obtained as a gBlock (IDT). The initial validation constructs were constructed via PCR and restriction cloning; genes were amplified with sticky ends from *B. burgdorferi* pBSV2 constructs and cloned using PstI and HindIII into pETN. Sequences were then sequence confirmed using Sanger sequencing. After initial validation using CspA, ElpQ, and INPN, other constructs were generated through Twist Biosciences to the same specifications.

To generate recombinant Elp homolog constructs free of tags DNA fragments were codon optimized for *Escherichia coli* and synthetically produced using the Integrated DNA Technologies gBlock Gene Fragment service with 5′BamHI and 3′NotI restriction sites appended. Each restriction enzyme-digested insert was ligated into pT7HMT and transformed into *E. coli* DH5α, using previously established protocols. (Garcia *et al*., 2016; Garrigues *et al*., 2022); (Geisbrecht *et al*., 2006; Thomas *et al*., 2024) Transformants were selected on LB agar plates containing kanamycin (50 μg/mL), and following plasmid isolation, sequences were confirmed (Eurofins).

### Optimization of conditions for B. burgdorferi lipoproteins on the surface of E. coli surface display

We optimized experimental conditions for surface display. By comparing various growth conditions, we found that strains were able to grow and surface express well when grown without induction overnight at 37°C. Those strains are grown for 15 hr at 37°C and used immediately. Strains expressed at variable levels under the derepression growth condition. As a result, some strains were grown to ∼0.6 OD_600_ at 37°C and induced using 0.4 mM IPTG for 18 hr at room temperature.

### Far western blotting

Strains were grown overnight (using the derepression growth condition) as previously described (Probert and Johnson, 1998). Cells were washed, resuspended with Laemmli+5% BME, and boiled as previously described. SDS-PAGE and PVDF membrane transfer was performed as previously described. PVDF membranes were incubated with 6M guanidine-HCl to denature proteins, followed by decreasing concentrations of guanidine-HCl to renature proteins. 6M guanidine-HCl was diluted in protein-binding buffer [20 mM Tris (pH 7.5), 0.1M sodium chloride, 1 mM EDTA, 1 mM DTT, 10% (v/v) glycerol, 0.1% (v/v) Tween-20, 5% (w/v) nonfat dry milk].

Proteins were incubated overnight with 2 µg/mL of the ligand (in protein-binding buffer), washed, and treated with either mouse anti-FH (Santa Cruz Biotechnology sc-53067) (1:50 dilution in Bio-Rad EveryBlot) or mouse anti-C1s (R&D Systems MAB2060), followed by anti-mouse HRP antibody (Promega W4021).

*E. coli flow cytometry:* 1×10^8^ cells per condition were collected from uninduced cultures of *E. coli* expressing a given protein and washed to remove LB. Cells were blocked for 2 hr with 3% BSA and incubated overnight with 1:500 dilution of rabbit anti-His antibody (Sigma-Aldrich SAB5600227). Cells were washed again and incubated for 1 hr with a 1:1000 dilution of either anti-rabbit FITC (Abcam 6717) or anti-rabbit Alexa Fluor 488 antibody (Invitrogen A11008). Cells were fixed with 4% formaldehyde and 30000 events were analyzed on a BD Accuri™ C6 Flow Cytometer. As negative controls, cells were compared to cells from the same cultures not treated with the secondary antibody.

### E. coli fluorescence microscopy

1×10^8^ cells per condition were collected and treated as previously described for flow cytometry. Cells were diluted in 5 mL and 10 µL were added to coverslips. Cells were fixed and observed at 100x magnification using an Echo Revolution through the FITC and Trans channels.

### Uninduced E. coli whole cell ELISA

Whole cell ELISA was performed as previously described (Pereira *et al*., 2022) with slight modifications. 8×10^6^ cells were collected to add 1×10^6^ cells in quadruplicate to C1 and BSA-coated wells. PBS was added to coated wells as a control in order to assess background. Cells were centrifuged to encourage cell-well contact, and allowed to incubate for 1 hr. Afterwards, non-adherent cells were washed away, and wells were incubated with HRP anti-His_6_ antibody. Binding was detected using TMB substrate and absorbance measured at OD_450_. The background level of signal from PBS wells was subtracted from OD_450_. The binding of a strain to its specific ligand can be compared directly to its binding to BSA because these wells contain the same strain from the same culture, expressing proteins at the same level.

Significance between specific and non-specific binding was determined via Student’s t-test. C1(A098) was obtained through Complementtech.

### Induced E. coli whole cell ELISA

Our screen was performed using the induced growth condition and the whole cell ELISA modification of 1×10^8^ cells per well. This is based on the relatively successful surface expression of Elps using the induced growth condition, as well as the knowledge that higher levels of surface expression did not always correlate with better detection of true binding interactions in one Elp-ECM component interaction (Fig. S5). 8×10^8^ cells were collected to add in quadruplicate to specific ligand-coated and BSA-coated wells of 96-well round-bottom ELISA plates (Nunc Maxisorp). C1r (A102) and C1s (A104) were obtained through Complementtech.

Perlecan was obtained through Sigma-Aldrich (H4777). PBS was added to coated wells as previously described. Cells were not centrifuged to accommodate the larger number of cells, and allowed to incubate for 2 hrs. Afterwards non-adherent cells were washed away as before, and adherent cells quantified as before.

### Protein expression and purification of recombinant proteins

Recombinant proteins used in recombinant protein ELISA were prepared as follows. Elp homolog sequences lacking the N-terminal signal sequence were ordered in a PGEX-4T2 background (Twist Bio) resulting in an N-terminally tagged GST recombinant protein. All constructs were confirmed through whole-plasmid sequencing prior to shipping. E. coli strain BL21 DE3 (NEB) were transformed with the vectors and positive colonies were selected from LB Agar plates supplemented with 100µg/mL of Ampicillin. Selected colonies were grown overnight at 37°C and transferred to fresh LB media the next day. Cultures were allowed to grow at 37°C for 2-5 hr until OD≈0.5-0.6. Next, 1 mM IPTG was added to the cultures. Cultures were induced overnight at 28°C and pelleted the following day before being frozen at -80C for future use. To purify the proteins of interest pelleted cells were resuspended in PBS and lysed 2-3x times at 15,000 psi using an LM20 Microfluidizer (Microfluidics). The lysed cells were then clarified at 12,000 rpm for 15 min at 4°C and the supernatant was collected. Supernatant was applied to Econofit Profinity GST Column (Bio-Rad) and bound GST-tagged protein was eluted from the column with elution buffer (30mM Reduced Glutathione, 100 mM Tris-HCL pH 8.0, 5 mM EDTA). Elution fractions with high 280nm values were pooled. Pooled fractions were concentrated, and buffer exchanged using Pierce Protein Concentrator PES (Thermo Scientific) before being validated via SDS-PAGE and quantified using a BCA assay (Thermo Fisher) according to manufacturer’s instructions. Protein was then aliquoted and stored at -80C for downstream assays.

Recombinant proteins for SPR and ELISA-based classical pathway inhibition assays were produced as follows. Elp homologs lacking affinity tags were produced in *E. coli* BL21(DE3) and purified using methods previously described.(Pereira *et al*., 2022; Garrigues *et al*., 2022; Thomas *et al*., 2024) Each construct was designed to remove to truncate the signal sequence and a disordered linker. Residues encoded in each construct were as follows: BG032 (residues 61-354), BG036 (residues 61-352), BG038 (residues 61-333), and BG044 (residues 61-368). Recombinant protein versions of Elp homologs and controls were produced in *E. coli*. These proteins were produced using PGEX-4T2 in BL21 (DE3). Proteins contained an N-terminal GST tag, the mature sequence of the lipoprotein, and a C-terminal His-Tag. Constructs were induced with IPTG, lysed through microfluidics, and purified using a Bio-Rad NGC chromatography system and Glutathione Sepharose columns Proteins were and screened via SDS-PAGE and quantified via BCA assay before being utilized in recombinant protein ELISA.

### Recombinant protein ELISA

ELISA was performed by coating wells as previously described, before incubating purified protein in PBS-T+3% BSA for 1.5 hr. 96-well round-bottom ELISA plates (Nunc Maxisorp) were used. Non-adherent protein was washed away and a 1:5000 dilution of rabbit anti-GST (Abcam 3445) was incubated for 1 hr. Wells were washed, incubated with a 1:8000 dilution of anti-rabbit HRP (Promega W4011) for 1 hr, washed, and incubated with TMB. Binding was quantified as previously described.

### Surface Plasmon Resonance (SPR)

SPR-binding assays were conducted as previously outlined. (Geisbrecht *et al*., 2006; Garcia *et al*., 2016; Pereira *et al*., 2022; Garrigues *et al*., 2022; Thomas *et al*., 2024) Elp homologs were immobilized onto a CMD200 sensor chip (XanTec bioanalytics) via amine coupling, using concentrations of 10 μg/mL in 10 mM sodium acetate (pH 5.0; pH 4.0 for BG038). The final immobilization densities, quantified in resonance units (RU), were as follows: BG036 (2298.9 RU), BG044 (1680.7 RU), BG032 (2867.8 RU), and BG038 (2898.2 RU). All assays were performed in a running buffer of HBS-T with Ca^2+^ (10 mM HEPES, pH 7.3, 140 mM NaCl, 0.005% [v/v] Tween-20, 5 mM CaCl_2_) at a 30 μl/min flow rate. Prior to and following each analyte injection, analytes were equilibrated in matching running buffer. Surfaces were regenerated to baseline with three 60-s injections of 2 M NaCl. For assessing interactions of Elp homologs with C1s or C1r, multicycle experiments were performed using active C1s or C1r (Complement Technology). An injection series (0, 0.8, 1.6, 3.1, 6.3, 12.5, 25, 50, 100, and 200 nM) was applied over an association time of 120 s, followed by a dissociation time of 180 s. Each injection series was conducted in triplicate and fit to a kinetic 1:1 binding model (Langmuir). Equilibrium dissociation constants (K_D_) were calculated from the resulting rate constants using the formula: K_D_= k_d_/k_a_ where k_d_ is the dissociation rate constant and k_a_ is the association rate constant. Fits of the reference- and background-corrected sensograms were determined using Biacore T200 Evaluation Software (v 3.2, Cytiva). Standard deviations were calculated based on triplicate injection series (n = 3).

### ELISA-based classical pathway inhibition assay

CP ELISA assays were conducted following previous protocols. (Garrigues *et al*., 2022; Thomas *et al*., 2024)Human IgM (3 μg/mL) (MP Biomedical) in coating buffer (100 mM Na_2_CO_3_/NaHCO_3_, pH 9.6) was immobilized on a high-binding ELISA plate (Greiner Bio-One). To assess dose-dependent BG inhibition, we used a 12-point, two-fold dilution: BG036 61-352 (4.4– 0.002 μM), BG044 (4.4–0.002 μM), BG032 (8.8–0.004 μM), and BG038 (8.8–0.004 μM).

Normal human serum (2%, Innovative Research) and the respective inhibitor in CP buffer (10 mM HEPES, pH 7.3, 0.1% [w/v] gelatin, 140 mM NaCl, 2 mM CaCl_2_, 0.5 mM MgCl_2_) were then added. All assays were performed in triplicate and normalized against positive (100%, no inhibitor) and negative controls (0%, no normal human serum). Nonlinear regression was used to fit the inhibition curves to a normalized variable slope model in GraphPad Prism (v 10.2.2, GraphPad Software, Inc).

### Borrelia flow cytometry

2×10^7^ cells per condition were collected from uninduced cultures of *E. coli* expressing a given protein and washed to remove BSK-II. Cells were blocked for 2 hr with 3% BSA and incubated for 1 hr with 1:500 dilution of rabbit anti-HA antibody (Abcam 20084). Cells were washed again and incubated for 1 hr with a 1:1000 dilution of anti-rabbit Alexa Fluor 488 antibody (Invitrogen A11008). Cells were fixed with 4% formaldehyde and 5000 events were analyzed on a BD Accuri™ C6 Flow Cytometer. As negative control, cells were compared to cells from a vector control culture treated with the secondary antibody.

### Borrelia whole cell ELISA

*Borrelia* whole cell ELISA was performed as previously described (Pereira *et al*., 2022) with slight modifications. Plates were coated with perlecan and BSA overnight at 4°C. Unbound ligand was washed away with 3X PBS-T washes and blocked with Ultrablock (Bio-Rad) for 2 hr. Wells were washed again 3X with PBS-T. 1.6×10^7^ cells were collected to add 4×10^6^ cells in quadruplicate to perlecan and BSA-coated wells. Cells were diluted in incomplete BSK-II media. Incomplete BSK-II was added to coated wells as a control in order to assess background. Cells were centrifuged to encourage cell-well contact, and allowed to incubate for 2 hr. Afterwards, non-adherent cells were washed away and adherent cells fixed for 20 min with 100 µL 10% buffered formalin. Formalin was removed and wells allowed to dry overnight. Wells were washed 1X with PBS-T and blocked with 5% milk for 1 hr. Wells were incubated with 1:10000 rabbit anti-HA in PBS-T + 5% milk and 1:2500 anti-rabbit HRP in PBS-T + 5% milk for 1 hr each. Binding was detected using TMB substrate and absorbance measured at OD_450_. The background level of signal from PBS wells was subtracted from OD_450_. Binding of a strain to perlecan was compared to its binding to BSA because these wells contain the same strain from the same culture, expressing proteins at the same level. Significance between specific and non-specific binding was determined via Student’s t-test.

## Acknowledgments

We thank Yun Tian and Michael Foster for valuable and practical advice and discussion. Support for this work was provided by Public Health Service grants K99/R)0AI148604 (to J.E.L.) and R01-AI146930 from the National Institute of Allergy and Infectious Diseases, National Institutes of Health (to B. L. G.).

## Author Contributions

**Nathan Hill:** Conceptualization, Formal Analysis, Investigation, Methodology, Project Administration, Visualization, Writing-Original Draft Preparation, Writing-Review and Editing; **Lara M. Matulina:** Formal Analysis, Investigation, Methodology, Validation, Visualization; **Cameron MacIntyre:** Formal Analysis, Investigation, Methodology, Validation, Writing-Review and Editing; **M. Amine Hassani:** Investigation, Validation, Writing-Original Draft Preparation, Writing-Review and Editing; **Sheila Thomas:** Investigation, Methodology, Writing-Review and Editing; **Matteo Luban:** Formal Analysis, Investigation, Methodology, Validation; **Isabelle Ward:** Investigation, Validation; **Amina Abdalla:** Investigation, Validation, Writing-Review and Editing; **John M. Leong:** Methodology, Supervision, Resources, Writing – Review & Editing; **Brandon L. Garcia:** Funding Acquisition, Methodology, Supervision, Resources, Writing-Review and Editing; **Jacob E. Lemieux:** Conceptualization, Funding Acquisition, Project Administration, Resources, Software, Supervision, Writing – Review & Editing

## References

Akins Darrin R., Caimano Melissa J., Yang Xiaofeng, Cerna Felix, Norgard Michael V., and Radolf Justin D. (1999) Molecular and Evolutionary Analysis ofBorrelia burgdorferi 297 Circular Plasmid-Encoded Lipoproteins with OspE- and OspF-Like Leader Peptides. Infect Immun 67: 1526–1532.

Akther, S., Mongodin, E.F., Morgan, R.D., Di, L., Yang, X., Golovchenko, M., et al. (2024) Natural selection and recombination at host-interacting lipoprotein loci drive genome diversification of Lyme disease and related bacteria. MBio e0174924.

Alvarez-Dominguez, C., and Vazquez-Boland, J.A. Host cell heparan sulfate proteoglycans mediate attachment and entry of Listeria monocytogenes, and the listerial surface protein ActA is involved in heparan sulfate …. Infection https://journals.asm.org/doi/abs/10.1128/iai.65.1.78-88.1997.

Antonara, S., Ristow, L., and Coburn, J. (2011) Adhesion mechanisms of Borrelia burgdorferi. Adv Exp Med Biol 715: 35–49.

Balmelli, T., and Piffaretti, J.C. (1995) Association between different clinical manifestations of Lyme disease and different species of Borrelia burgdorferi sensu lato. Res Microbiol 146: 329– 340.

Barbour, A.G. (1984) Isolation and cultivation of Lyme disease spirochetes. Yale J Biol Med 57: 521–525.

Bouchard, C., Leonard, E., Koffi, J.K., Pelcat, Y., Peregrine, A., Chilton, N., et al. (2015) The increasing risk of Lyme disease in Canada. Can Vet J 56: 693–699.

Brissette, C.A., Verma, A., Bowman, A., Cooley, A.E., and Stevenson, B. (2009) The Borrelia burgdorferi outer-surface protein ErpX binds mammalian laminin. Microbiology 155: 863–872.

Caine, J.A., Lin, Y.-P., Kessler, J.R., Sato, H., Leong, J.M., and Coburn, J. (2017) Borrelia burgdorferi outer surface protein C (OspC) binds complement component C4b and confers bloodstream survival. Cell Microbiol 19 10.1111/cmi.12786.

Casjens, S. (1999) Evolution of the linear DNA replicons of the Borrelia spirochetes. Curr Opin Microbiol 2: 529–534.

Casjens, S., Palmer, N., Vugt, R. van, Huang, W.M., Stevenson, B., Rosa, P., et al. (2000) A bacterial genome in flux: the twelve linear and nine circular extrachromosomal DNAs in an infectious isolate of the Lyme disease spirochete Borrelia burgdorferi. Mol Microbiol 35: 490– 516.

Casjens, S.R., Gilcrease, E.B., Vujadinovic, M., Mongodin, E.F., Luft, B.J., Schutzer, S.E., et al. (2017) Plasmid diversity and phylogenetic consistency in the Lyme disease agent Borrelia burgdorferi. BMC Genomics 18: 165.

Chen, L.D., and Hazlett, L.D. (2000) Perlecan in the basement membrane of corneal epithelium serves as a site for P. aeruginosa binding. Curr Eye Res 20: 260–267.

Dam, A.P. van, Kuiper, H., Vos, K., Widjojokusumo, A., Jongh, B.M. de, Spanjaard, L., et al. (1993) Different genospecies of Borrelia burgdorferi are associated with distinct clinical manifestations of Lyme borreliosis. Clin Infect Dis 17: 708–717.

Dowdell, A.S., Murphy, M.D., Azodi, C., Swanson, S.K., Florens, L., Chen, S., and Zückert, W.R. (2017) Comprehensive Spatial Analysis of the Borrelia burgdorferi Lipoproteome Reveals a Compartmentalization Bias toward the Bacterial Surface. J Bacteriol 199 10.1128/JB.00658-16.

Dykhuizen, D.E., Brisson, D., Sandigursky, S., Wormser, G.P., Nowakowski, J., Nadelman, R.B., and Schwartz, I. (2008) The Propensity of Different Borrelia burgdorferi sensu stricto Genotypes to Cause Disseminated Infections in Humans. Am J Trop Med Hyg 78: 806–810.

Fan, L.-H., Liu, N., Yu, M.-R., Yang, S.-T., and Chen, H.-L. (2011) Cell surface display of carbonic anhydrase on Escherichia coli using ice nucleation protein for COLJ sequestration. Biotechnol Bioeng 108: 2853–2864.

Fischer, J.R., LeBlanc, K.T., and Leong, J.M. (2006) Fibronectin binding protein BBK32 of the Lyme disease spirochete promotes bacterial attachment to glycosaminoglycans. Infect Immun 74: 435–441.

Fraser, C.M., Casjens, S., Huang, W.M., Sutton, G.G., Clayton, R., Lathigra, R., et al. (1997) Genomic sequence of a Lyme disease spirochaete, Borrelia burgdorferi. Nature 390: 580–586.

Garcia, B.L., Zhi, H., Wager, B., Höök, M., and Skare, J.T. (2016) Borrelia burgdorferi BBK32 Inhibits the Classical Pathway by Blocking Activation of the C1 Complement Complex. PLoS Pathog 12: e1005404.

Garrigues, R.J., Thomas, S., Leong, J.M., and Garcia, B.L. (2022) Outer surface lipoproteins from the Lyme disease spirochete exploit the molecular switch mechanism of the complement protease C1s. J Biol Chem 298: 102557.

Geisbrecht, B.V., Bouyain, S., and Pop, M. (2006) An optimized system for expression and purification of secreted bacterial proteins. Protein Expr Purif 46: 23–32.

Grillon, A., Scherlinger, M., Boyer, P.-H., De Martino, S., Perdriger, A., Blasquez, A., et al. (2018) Characteristics and clinical outcomes after treatment of a national cohort of PCR-positive Lyme arthritis. Semin Arthritis Rheum 10.1016/j.semarthrit.2018.09.007.

Grimm, D., Elias, A.F., Tilly, K., and Rosa, P.A. (2003) Plasmid stability during in vitro propagation of Borrelia burgdorferi assessed at a clonal level. Infect Immun 71: 3138–3145.

Grossman, T.H., Kawasaki, E.S., Punreddy, S.R., and Osburne, M.S. (1998) Spontaneous cAMP-dependent derepression of gene expression in stationary phase plays a role in recombinant expression instability. Gene 209: 95–103.

Haake, D.A. (2000) Spirochaetal lipoproteins and pathogenesis. Microbiology 146 ( **Pt 7**): 1491–1504.

Hayes, A.J., Farrugia, B.L., Biose, I.J., Bix, G.J., and Melrose, J. (2022) Perlecan, A Multi- Functional, Cell-Instructive, Matrix-Stabilizing Proteoglycan With Roles in Tissue Development Has Relevance to Connective Tissue Repair and Regeneration. Frontiers in Cell and Developmental Biology 10 https://www.frontiersin.org/journals/cell-and-developmental-biology/articles/10.3389/fcell.2022.856261.

Jones, K.L., Glickstein, L.J., Damle, N., Sikand, V.K., McHugh, G., and Steere, A.C. (2006) Borrelia burgdorferi genetic markers and disseminated disease in patients with early Lyme disease. J Clin Microbiol 44: 4407–4413.

Jones, K.L., McHugh, G.A., Glickstein, L.J., and Steere, A.C. (2009) Analysis of Borrelia burgdorferi genotypes in patients with Lyme arthritis: High frequency of ribosomal RNA intergenic spacer type 1 strains in antibiotic-refractory arthritis. Arthritis Rheum 60: 2174–2182.

Jung, H.C., Park, J.H., Park, S.H., Lebeault, J.M., and Pan, J.G. (1998) Expression of carboxymethylcellulase on the surface of Escherichia coli using Pseudomonas syringae ice nucleation protein. Enzyme Microb Technol 22: 348–354.

Kraiczy, P., Hellwage, J., Skerka, C., Becker, H., Kirschfink, M., Simon, M.M., et al. (2004) Complement resistance of Borrelia burgdorferi correlates with the expression of BbCRASP-1, a novel linear plasmid-encoded surface protein that interacts with human factor H and FHL-1 and is unrelated to Erp proteins. J Biol Chem 279: 2421–2429.

Kraiczy, P., Skerka, C., Brade, V., and Zipfel, P.F. (2001) Further characterization of complement regulator-acquiring surface proteins of Borrelia burgdorferi. Infect Immun 69: 7800– 7809.

Kugeler, K.J., Schwartz, A.M., Delorey, M.J., Mead, P.S., and Hinckley, A.F. (2021) Estimating the Frequency of Lyme Disease Diagnoses, United States, 2010-2018. Emerg Infect Dis 27: 616–619.

Lantos, P.M., Nigrovic, L.E., Auwaerter, P.G., Fowler, V.G., Ruffin, F., Brinkerhoff, R.J., et al. (2015) Geographic Expansion of Lyme Disease in the Southeastern United States, 2000–2014. Open Forum Infect Dis 2: ofv143.

Lei, S.P., Lin, H.C., Wang, S.S., Callaway, J., and Wilcox, G. (1987) Characterization of the Erwinia carotovora pelB gene and its product pectate lyase. J Bacteriol 169: 4379–4383.

Lemieux, J.E. (2024) Analysis of the Borreliaceae pangenome reveals a distinct genomic architecture conserved across phylogenetic scales. J Infect Dis 230: S51–S61.

Lemieux, J.E., Huang, W., Hill, N., Cerar, T., Freimark, L., Hernandez, S., et al. (2023) Whole genome sequencing of human Borrelia burgdorferi isolates reveals linked blocks of accessory genome elements located on plasmids and associated with human dissemination. PLoS Pathog 19: e1011243.

Lin, Y.-P., Bhowmick, R., Coburn, J., and Leong, J.M. (2015) Host cell heparan sulfate glycosaminoglycans are ligands for OspF-related proteins of the Lyme disease spirochete. Cell Microbiol 17: 1464–1476.

Lin, Y.-P., Diuk-Wasser, M.A., Stevenson, B., and Kraiczy, P. (2020a) Complement Evasion Contributes to Lyme Borreliae-Host Associations. Trends Parasitol 36: 634–645.

Lin, Y.-P., Tan, X., Caine, J.A., Castellanos, M., Chaconas, G., Coburn, J., and Leong, J.M. (2020b) Strain-specific joint invasion and colonization by Lyme disease spirochetes is promoted by outer surface protein C. PLoS Pathog 16: e1008516.

McDowell, J.V., Hovis, K.M., Zhang, H., Tran, E., Lankford, J., and Marconi, R.T. (2006) Evidence that the BBA68 protein (BbCRASP-1) of the Lyme disease spirochetes does not contribute to factor H-mediated immune evasion in humans and other animals. Infect Immun 74: 3030–3034.

Mead, P.S. (2015) Epidemiology of Lyme disease. Infect Dis Clin North Am 29: 187–210.

Norris, S.J., Howell, J.K., Garza, S.A., Ferdows, M.S., and Barbour, A.G. (1995) High- and low-infectivity phenotypes of clonal populations of in vitro-cultured Borrelia burgdorferi. Infect Immun 63: 2206–2212.

Pereira, M.J., Wager, B., Garrigues, R.J., Gerlach, E., Quinn, J.D., Dowdell, A.S., et al. (2022) Lipoproteome screening of the Lyme disease agent identifies inhibitors of antibody-mediated complement killing. Proc Natl Acad Sci U S A 119: e2117770119.

Probert, W.S., and Johnson, B.J. (1998) Identification of a 47 kDa fibronectin-binding protein expressed by Borrelia burgdorferi isolate B31. Mol Microbiol 30: 1003–1015.

Radolf, J.D., Strle, K., Lemieux, J.E., and Strle, F. (2021) Lyme Disease in Humans. Curr Issues Mol Biol 42: 333–384.

Robertson, K.E., Truong, C.D., Craciunescu, F.M., Yang, J.-H., Chiu, P.-L., Fromme, P., and Hansen, D.T. (2019) Membrane directed expression in Escherichia coli of BBA57 and other virulence factors from the Lyme disease agent Borrelia burgdorferi. Sci Rep 9: 17606.

Rudenko, N., Golovchenko, M., Grubhoffer, L., and Oliver, J.H., Jr (2011) Updates on Borrelia burgdorferi sensu lato complex with respect to public health. Ticks Tick Borne Dis 2: 123–128.

Samuels, D.S. (1995) Electrotransformation of the spirochete Borrelia burgdorferi. Methods Mol Biol 47: 253–259.

Seshu, J., Moy, B.E., and Ingle, T.M. (2021) Transformation of Borrelia burgdorferi. Curr Protoc 1: e61.

Skare, J.T., and Garcia, B.L. (2020) Complement Evasion by Lyme Disease Spirochetes. Trends Microbiol 28: 889–899.

Spillmann, D. (2001) Heparan sulfate: anchor for viral intruders? Biochimie 83: 811–817.

Steere, A.C., Strle, F., Wormser, G.P., Hu, L.T., Branda, J.A., Hovius, J.W.R., et al. (2016) Lyme borreliosis. Nat Rev Dis Primers 2: 16090.

Strle, K., Jones, K.L., Drouin, E.E., Li, X., and Steere, A.C. (2011) Borrelia burgdorferi RST1 (OspC type A) genotype is associated with greater inflammation and more severe Lyme disease. Am J Pathol 178: 2726–2739.

Tan, X., Petri, B., DeVinney, R., Jenne, C.N., and Chaconas, G. (2021) The Lyme disease spirochete can hijack the host immune system for extravasation from the microvasculature. Mol Microbiol 116: 498–515.

Thomas, S., Schulz, A.M., Leong, J.M., Zeczycki, T.N., and Garcia, B.L. (2024) The molecular determinants of classical pathway complement inhibition by OspEF-related proteins of Borrelia burgdorferi. J Biol Chem 300: 107236.

Wormser, G.P., Brisson, D., Liveris, D., Hanincová, K., Sandigursky, S., Nowakowski, J., et al. (2008) Borrelia burgdorferi genotype predicts the capacity for hematogenous dissemination during early Lyme disease. J Infect Dis 198: 1358–1364.

Wormser, G.P., Liveris, D., Nowakowski, J., Nadelman, R.B., Cavaliere, L.F., McKenna, D., et al. (1999) Association of specific subtypes of Borrelia burgdorferi with hematogenous dissemination in early Lyme disease. J Infect Dis 180: 720–725.

