## Supplementary Materials for "Heterologous Surface Display Reveals Conserved Complement Inhibition and Functional Diversification of *Borrelia burgdorferi* Elp Proteins"

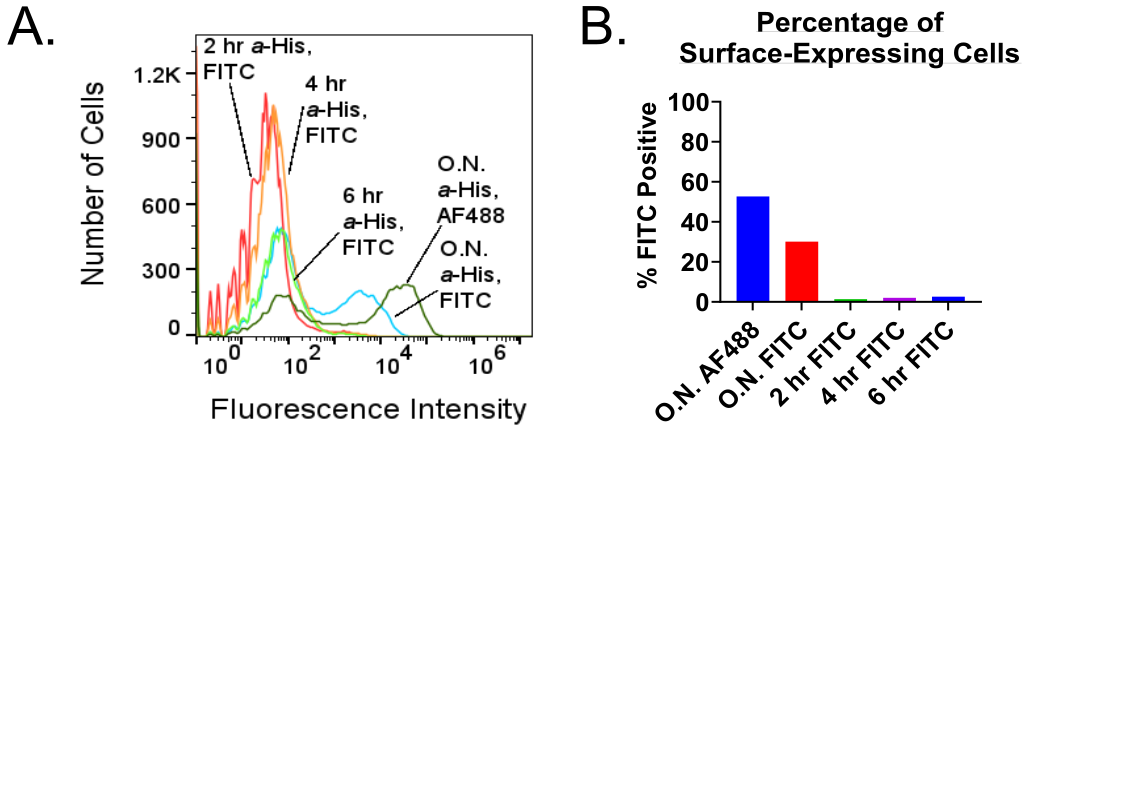

Supplemental Figure 1. **Optimization of growth and induction conditions for surface expression:** (A). CspA was grown overnight to stationary phase (OD≈1.5). Cells were harvested from this culture as well as cultures inoculated from the overnight culture (O.N.) at a 1:100 dilution and allowed to grow for 2 hr, 4 hr, and 6 hr. Cells were washed and treated with anti-His antibodies and either anti-rabbit FITC or anti-rabbit AF488 antibodies. Flow cytometry performed as described (Methods). (B). Percentage of FITC positivity for cells grown under the different conditions and treated with different antibodies. Positivity based on gating above background level of fluorescence for a control not treated with fluorophore-conjugated antibody.

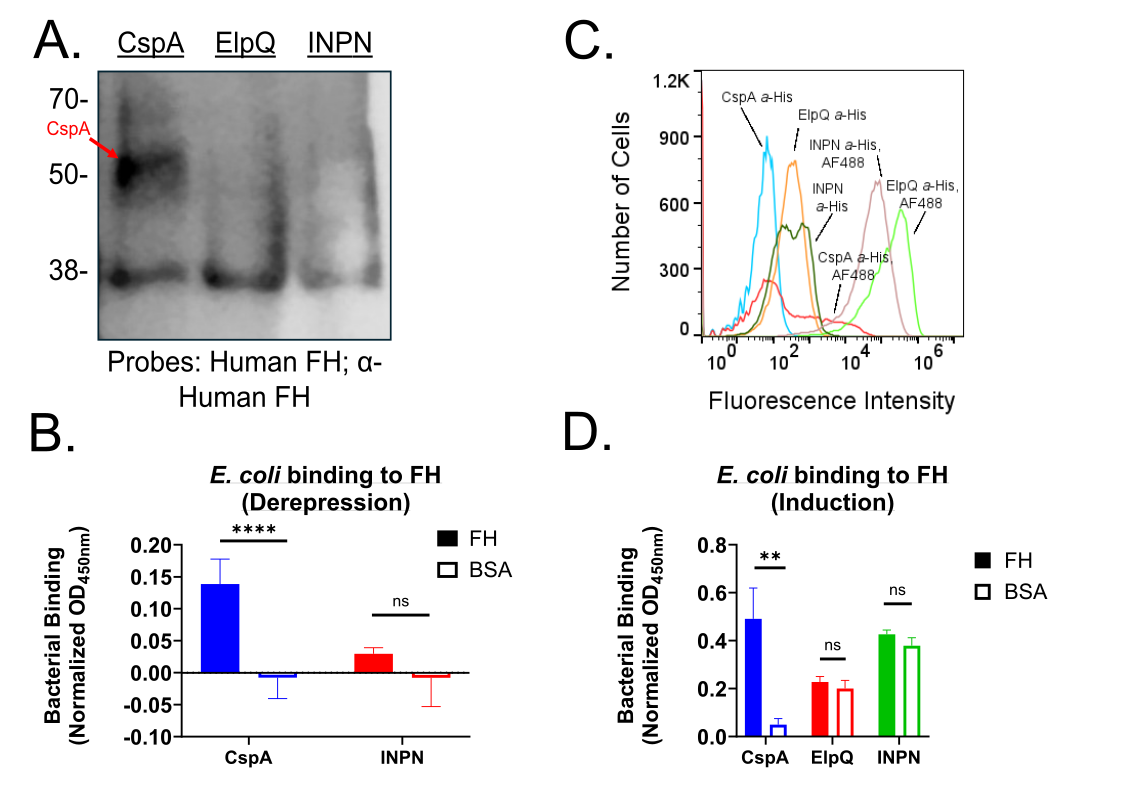
 Supplemental Figure 2. **Validation and optimization of CspA ligand-specific binding activity upon ectopic expression on the surface of *E. coli*:** (A). Far western blotting was performed as previously described. Protein-containing PVDF membranes were fully denatured and renatured through application and down-titration of a denaturant (guanidine-HCl). Proteins were incubated overnight with 2 µg/mL of FH, washed, and treated with mouse anti-FH antibody, followed by an HRP-conjugated antibody. (B). Cells were grown overnight under the derepression protein expression protocol as previously described. 1x10^6^ cells of a given strain were added to wells in quadruplicate, centrifuged to encourage contact with FH-coated and BSA-coated wells, and incubated for 1 hr. Binding was detected as previously described. Experiment was performed in duplicate (n=2). (C). Strains expressing CspA-INPN, ElpQ-INPN, and INPN were grown to ~0.6 OD_600_ and induced for 18 hr as previously described. 8x10^8^ cells were collected to add in quadruplicate to FH and BSA-coated wells. PBS was added to coated wells as previously described. Cells were allowed to incubate for 2 hrs. Afterwards, non-adherent cells were washed away, and wells were incubated with HRP anti-His_6_-tag antibody. Binding was detected using TMB substrate and absorbance measured at OD_450_. The background level of signal from PBS wells was subtracted from OD_450_. Significance between specific and non-specific binding was determined via Student’s t-test. CspA and ElpQ binding was assessed in at least triplicate experiments (n=4 and n=3, respectively). INPN binding was assessed in duplicate (n=2). (D). Strains expressing CspA, ElpQ, and INPN were grown under the induction growth condition and analyzed through flow cytometry as described (Methods). Fluorescence intensity of cells treated with rabbit anti-His antibody and either untreated or treated with anti-rabbit AF488 antibody are displayed.

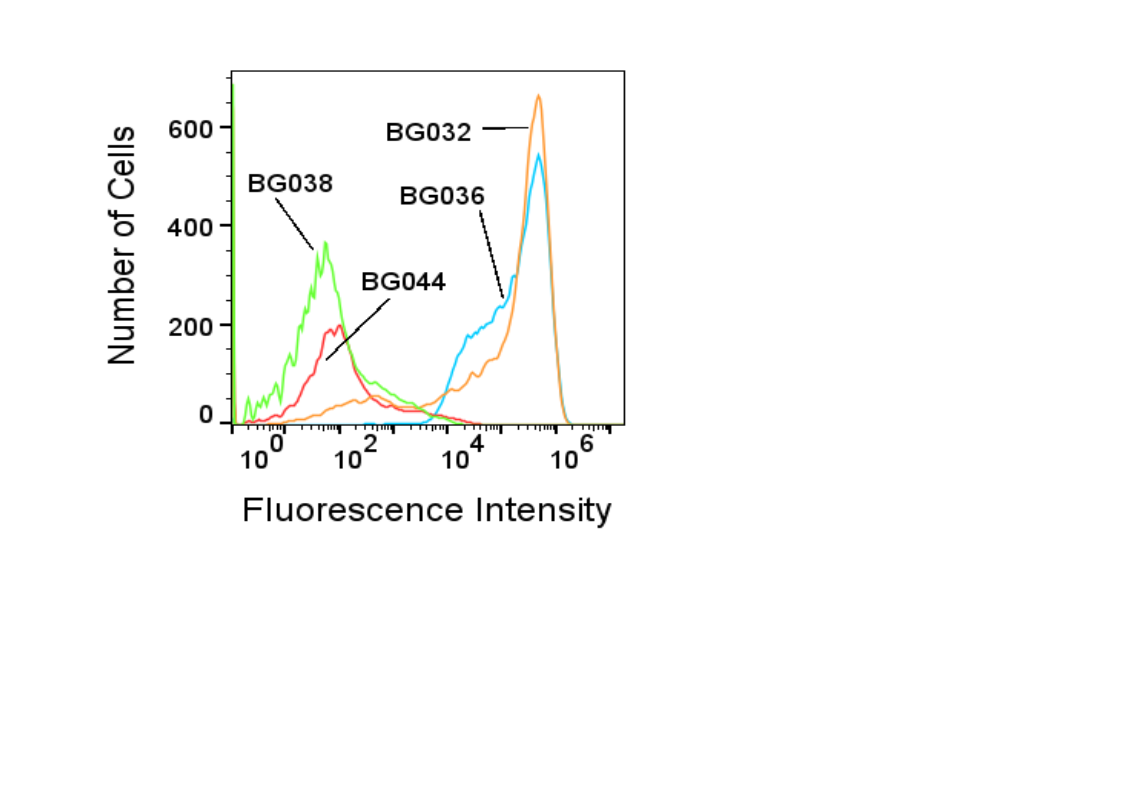

Supplemental Figure 3. **Induced cultures allow for surface display at meaningful levels for binding assessment:** Strains expressing BG036, BG044, BG032, and BG038 were grown under the induction growth condition and analyzed through flow cytometry as previously described. Fluorescence intensity of cells treated with both rabbit anti-His and anti-rabbit AF488 antibodies are displayed.

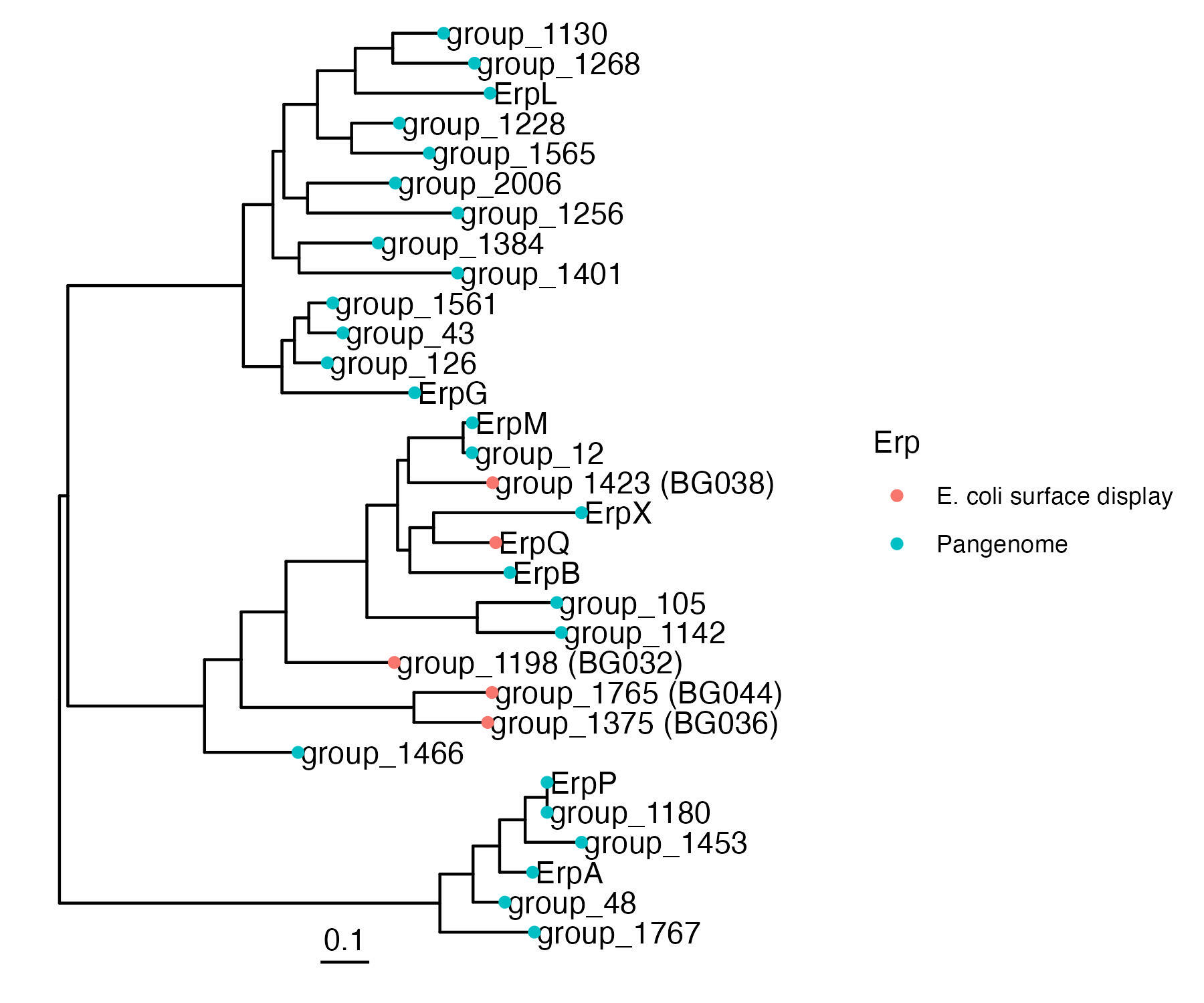

Supplementary Figure 4: **Erps and Elps from pangenome analysis:** Maximum likelihood tree of Erp sequences from *B. burgdorferi* sensu lato pangenome with B31 alleles noted. The homology groups studied are marked with an orange point.

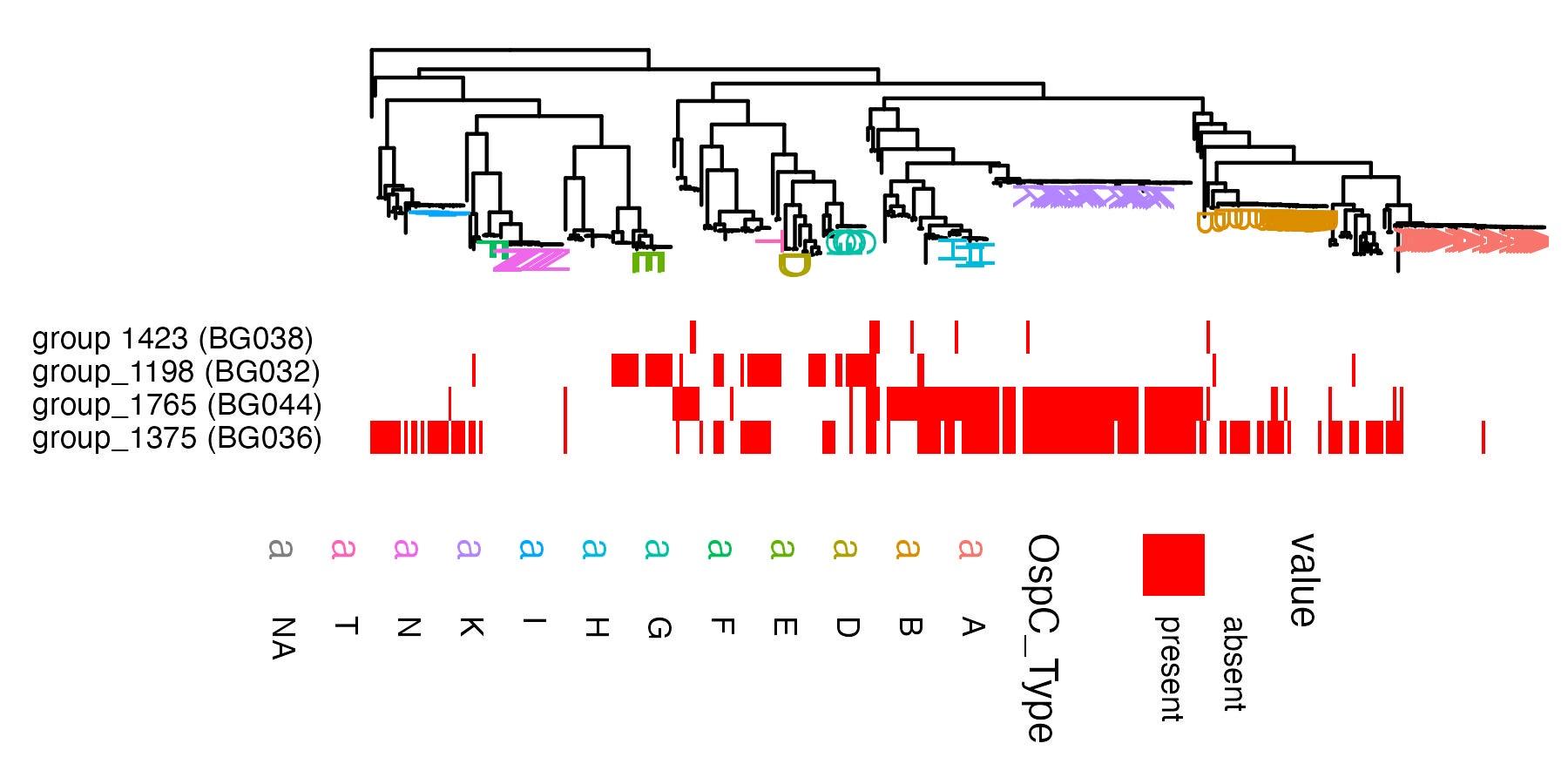

Supplementary Figure 5: **The phylogenetic distribution of Elp homologs studied:** A phylogenetic tree of *B. burgdorferi* sensu stricto isolates with OspC types marked. A matrix of presence / absence is plotted adjacent to tips. If an allele is present in a given strain, the matrix element is colored with a red bar.

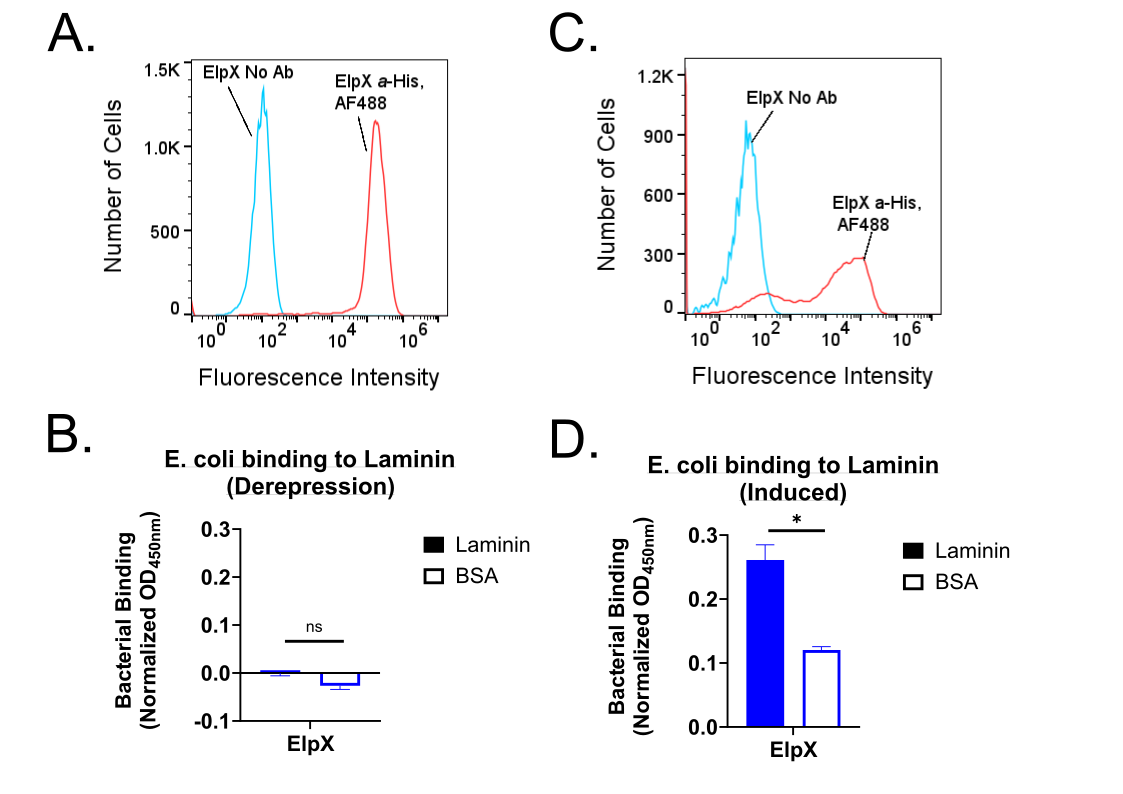

Supplemental Figure 6. **ElpX expressed at a moderate level performs better in a functional assay than ElpX expressed at a high level:** (A). An ElpX-expressing strain was grown under derepression growth conditions. Flow cytometry was performed as previously described. Fluorescence intensity of cells either untreated or treated with rabbit anti-His and anti-rabbit AF488 are displayed. (B). 1x10^6^ cells of ElpQ-expressing and ElpX-expressing strains were grown under derepression conditions. These cells were added to laminin-coated and BSA-coated wells in quadruplicate, centrifuged to encourage contact with wells, and incubated for 1 hr. Binding was detected as previously described. (C). 1x10^8^ cells of ElpQ-expressing and ElpX-expressing strains were grown under induction conditions. 8x10^8^ cells were collected from both strains to add in quadruplicate to FH and BSA-coated wells. PBS was added to coated wells as previously described. Cells were not centrifuged to accommodate the larger number of cells and allowed to incubate for 2 hrs. Binding was detected as previously described. (D). An ElpX-expressing strain was grown under induced growth conditions. Flow cytometry was performed as previously described. Fluorescence intensities of cells treated either untreated or treated with rabbit anti-His and anti-rabbit AF488 are displayed.

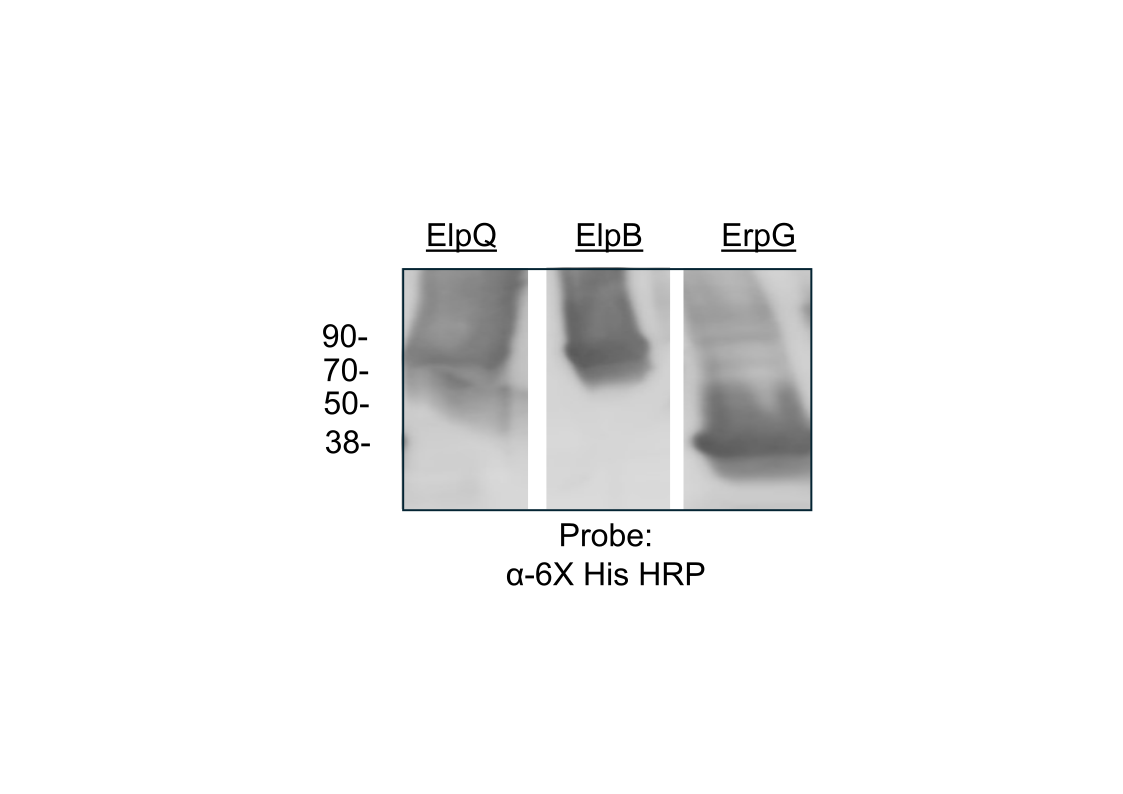

Supplementary Figure 7: **Western blotting shows protein production for ElpQ, ElpB, and ErpG strains grown under derepression growth conditions:** Strains expressing ElpQ-INPN, ElpB-INPN, and ErpG-INPN fusions were constructed and grown under derepression growth conditions. Western blotting was performed as previously described. All 3 strains produced bands around the expected size of their proteins. ElpQ was expected to produce a band around 62.6 kDa, ElpB was expected to produce a band around 66.5 kDa, and ErpG was expected to produce a band around 46.1 kDa.

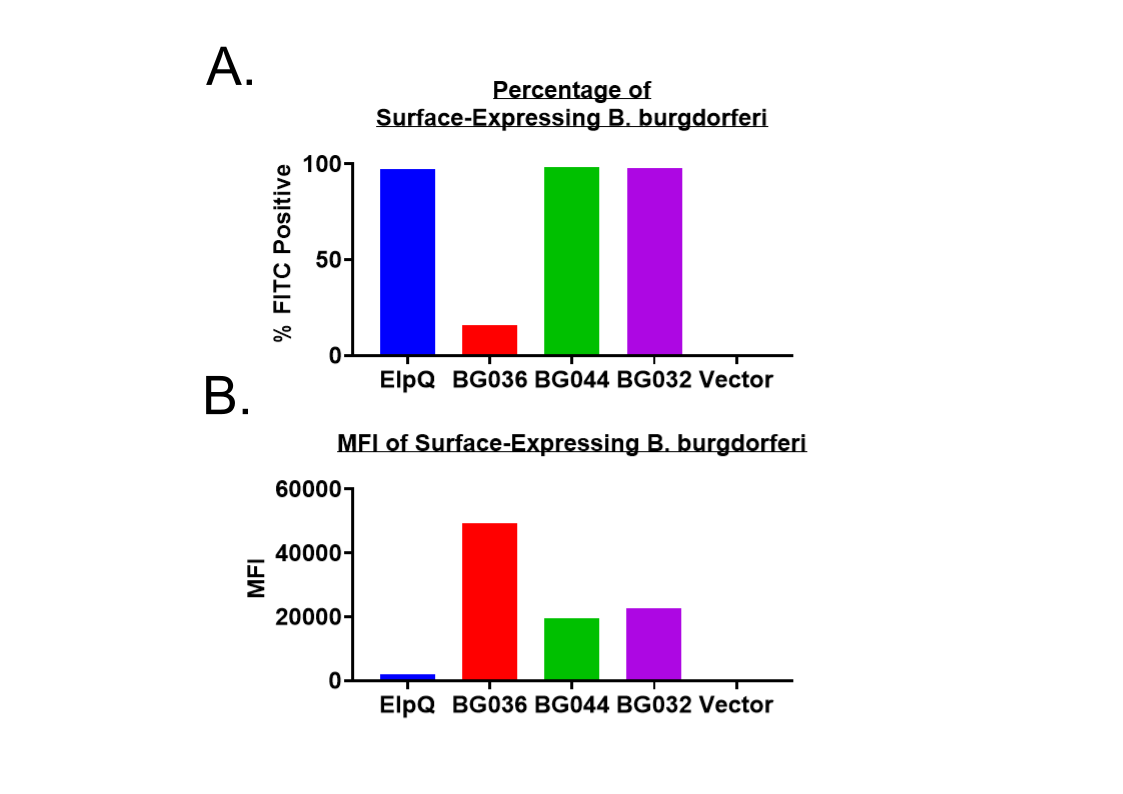

Supplementary Figure 8: **Surface expression of Elp homologs in *B. burgdorferi* is variable:** *B. burgdorferi* strains expressing ElpQ, BG036, BG044, BG032, and pBSV2 vector alone were grown to midlog. Cells were harvested, washed, and treated with a 1:500 of an anti-HA AF488 antibody. Flow cytometry was performed as described (Methods). Flow cytometry was performed. (A). The percentage of cells fluorescing at a level higher than the vector control was assessed. (B). The median fluorescence intensity of cells considered to be surface expressed was assessed.

**Supplemental Table 1:**

| **Name** | **Best Named Match BLASTx against NR (%)** | **Best Named Match in B31 (ORF, %)** | **Previously Known Binding Interactions for Best B31 Match** | **Strains/Species** | **Accession** |
| --- | --- | --- | --- | --- | --- |
| CspA | CspA B31 (100%) | CspA (BBA68, 100%) | FH, FHL-1 (Kraiczy *et al.*, 2004) | B31 | WP_106021774.1 |
| ElpQ | ElpQ B31 (100%) | ElpQ (100%, BBN39) | C1, C1r, C1s (Pereira *et al.*, 2022) | B31 | WP_010883887.1 |
| ElpX | ElpX B31 (100%) | ElpX (100%, BBQ47) | Laminin (Brissette *et al.*, 2009) | B31 | WP_010883925.1 |
| ElpB | ElpB B31 (100%) | ElpB (BBP39, 100%) | C1, C1r, C1s (Pereira *et al.*, 2022) | B31 | WP_010883724.1 |
| ErpG | ErpG B31 (100%) | ErpG (BBS41, 100%) | Heparan Sulfate, Heparin (Lin *et al.*, 2015) | B31 | WP_010883762.1 |
| BG036 | Erp43 WI91-23 (100%) | ElpB (BBP39, 59.1%) | C1, C1r, C1s (Pereira *et al.*, 2022) | WI91-23, B408, B331 | WP_012665139.1 |
| BG044 | Erp45 156a (100%) | ElpM(BBO40, 59.4%) | None | B379, 297, 156a | WP_012621421.1 |
| BG032 | Erp26 72a (100%) | ElpM (BBO40, 68.8%) | None | 72a, 118a | WP_012644033.1 |
| BG038 | ErpM JD1 (100%) | ElpB (BBP39, 68.8%) | C1, C1r, C1s (Pereira *et al.*, 2022) | JD1, 156a | WP_012621340.1 |

**Supplemental Table 2:**

| **Name** | **Vector** | **Size (kb)** | **Insert** | **Size (bp)** | **Reference** |
| --- | --- | --- | --- | --- | --- |
| pETN | pETN | 6.1 | NA | NA | 16 |
| CspA-pETN | pETN | 6.1 | CspA | 683 | 13 |
| ElpQ-pETN | pETN | 6.1 | ElpQ | 975 | 13 |
| ElpQ N-pETN | pETN | 6.1 | ElpQ (19-206) | 552 | 18 |
| ElpQ C-pETN | pETN | 6.1 | ElpQ (168-343) | 516 | 18 |
| ElpX-pETN | pETN | 6.1 | ElpX | 975 | 13 |
| ElpB-pETN | pETN | 6.1 | ElpB | 1107 | 13 |
| BG036-pETN | pETN | 6.1 | BG036 | 996 | This study |
| BG044-pETN | pETN | 6.1 | BG044 | 1044 | This study |
| BG032-pETN | pETN | 6.1 | BG032 | 1002 | This study |
| BG038-pETN | pETN | 6.1 | BG038 | 945 | This study |

**Supplemental Table 3: C1s recombinant protein ELISA binding**

| **Protein** | **K_D_ (μM)** | **95% C.I. (μM)** |
| --- | --- | --- |
| BG036 | 0.004512 | 0 to 0.01494 |
| BG044 | 0.2302 | 0.03117 to 1.652 |
| BG032 | 0.1988 | 0.06428 to 0.6294 |
| BG038 | 0.1149 | 0.05482 to 0.2394 |

**Supplemental Table 4: C1r recombinant protein ELISA binding**

| **Protein** | **K_D_ (μM)** | **95% C.I. (μM)** |
| --- | --- | --- |
| BG036 | 0.5659 | 0.3184 to 1.034 |
| BG044 | N.B | NA |
| BG032 | 5.498 | 3.091 to 14.42 |
| BG038 | 4.003 | 2.279 to 9.360 |

**Supplemental Table 5: SPR binding parameters**

| **Protein** | **k_a_ (1/Ms)** | **k_d_ (1/s)** | **K_D, kin_ (M)** | **R_max_ (RU)** |
| --- | --- | --- | --- | --- |
| BG036 | 3.22 x 10^5^  (1.96 x 10^3^) | 3.01 x 10^-3^  (4.15 x 10^-5^) | 9.35 x 10^-9^  (1.69 x 10^-10^) | 906  (1.78) |
| BG044 | 8.49 x 10^5^  (54.7 x 10^3^) | 3.23 x 10^-2^  (8.29 x 10^-4^) | 3.82 x 10^-8^  (3.29 x 10^-9^) | 141  (3.98) |
| BG032 | 2.47 x 10^4^  (6.93 x 10^3^) | 7.21 x 10^-3^  (0.481 x 10^-3^) | 3.05 x 10^-7^  (7.13 x 10^-8^) | 174  (35.6) |
| BG038 | 1.10 x 10^5^  (7.23 x 10^3^) | 8.26 x 10^-3^  (6.79 x 10^-5^) | 7.56 x 10^-8^  (5.41 x 10^-9^) | 129  (0.115) |

Association rate constants are depicted as *k*_a_ and dissociation rate constants as *k*_d_. Equilibrium dissociation constants (*K*_D_) were calculated by performing kinetic analysis (*K*_D, kin_) for each interaction, using a 1:1 binding model. Values in parentheses indicate standard deviation (Methods).

**Supplemental Table 6: Classical Pathway ELISA potencies**

| **Protein** | **IC50 (nM)** | **95% CI (nM)** |
| --- | --- | --- |
| BG036 | 89.8 | 72.7 to 111 |
| BG044 | 139.7 | 125 to 157 |
| BG032 | 2595 | 1956 to 3582 |
| BG038 | 1544 | 1289 to 1859 |

IC50 values were obtained by nonlinear regression analysis using a four-parameter variable slope fit and constraining the *bottom* and *top* values to 0 and 100, respectively. The 95% confidence interval (95% CI) is shown.

**Supplemental Table 7: Perlecan recombinant protein ELISA binding**

| **Protein** | **K_D_ (μM)** |
| --- | --- |
| ElpQ | 7.031 |
| BG036 | 7.744 |
| BG044 | 11.49 |
| BG032 | N.B. |

**References:**

Akins Darrin R., Caimano Melissa J., Yang Xiaofeng, Cerna Felix, Norgard Michael V., and Radolf Justin D. (1999) Molecular and Evolutionary Analysis ofBorrelia burgdorferi 297 Circular Plasmid-Encoded Lipoproteins with OspE- and OspF-Like Leader Peptides. *Infect Immun* **67**: 1526–1532.
